# Polycystin-1 loss of function increases susceptibility to atrial fibrillation through impaired DNA damage response

**DOI:** 10.1101/2024.07.08.602618

**Authors:** Troy Hendrickson, Abigail Giese, Matthew Fiedler, William Perez, Ernesto Reyes-Sanchez, Monserrat Reyes-Lozano, Sufen Wang, Leslye Venegas-Zamora, Vincent Provasek, Aschraf El-Essawi, Ingo Breitenbach, Funsho Fakuade, Ingo Kutschka, Gabriele G. Schiattarella, Niels Voigt, Miguel Valderrábano, Francisco Altamirano

## Abstract

**Background:** The increasing prevalence of atrial fibrillation (AF) and chronic kidney diseases highlights the need for a deeper comprehension of the molecular mechanisms linking them. Mutations in *PKD1*, the gene encoding Polycystin-1 (PKD1 or PC1), account for 85% of autosomal dominant polycystic kidney disease (ADPKD) cases. This disease often includes cardiac complications such as AF. In cardiomyocytes, PC1 deletion reduces hypertrophic response to pressure overload but promotes baseline ventricular dysfunction, while deletion in fibroblasts ameliorates post-myocardial infarction fibrosis. Despite its known cardiac impact, the role of PC1 in atrial cardiomyocytes and arrhythmias is less understood. Here, we sought to investigate the role of PC1 in AF.

**Methods:** We used intracardiac programmed stimulation and optical mapping to evaluate AF inducibility in two mouse models, *Pkd1* R3277C, which recapitulates human ADPKD progression, and cardiomyocyte-specific *Pkd1* deletion, and their respective controls. Isolated adult mouse atrial cardiomyocytes, human iPSC-derived atrial cardiomyocytes (hiPSC-aCM), and HL-1 cells served as *in vitro* cellular models. Molecular mechanisms were evaluated using optical mapping and molecular and biochemical approaches.

**Results:** Loss-of-function PC1 mutations significantly increased AF susceptibility *in vivo* and facilitated local reentry in *ex vivo* left atrial appendages. Comprehensive *in vitro* experiments supported a direct effect of PC1 in atrial cardiomyocytes. PC1-deficient monolayers exhibited increased arrhythmic events, escalating into reentrant spiral waves post-tachypacing. Transcriptomics analysis revealed PC1-dependent regulation of DNA repair, with PC1 deficiency leading to increased DNA damage under stress. PARP1 inhibitors or nicotinamide riboside, which counteract DNA damage-related metabolic consequences, reduced *in vitro* arrhythmias PC1-deficient monolayers. Overexpression of the C-terminus of PC1 had the opposite effects in DNA repair genes, suggesting its regulatory effects in atrial cardiomyocytes through retinoblastoma/E2F. Analyses of human atrial tissue from non-ADPKD AF patients showed reduced levels of mature PC1, suggesting a broader relevance of impaired PC1 in AF.

**Conclusions:** Impaired PC1 increases *in vivo* AF inducibility under programmed electrical stimulation and promotes *in vitro* arrhythmias in hiPSC-aCM and HL-1 cells. Our findings indicate that PC1 protects against DNA damage to reduce AF susceptibility.

## Introduction

Atrial fibrillation (AF) – the most prevalent sustained adult arrhythmia – is a growing epidemic and a major healthcare and economic burden.^1^ AF consists of multiple rapid, uncoordinated propagation wavefronts that impair normal atrial contractility,^1,2^ leading to local stasis, thrombogenesis, heart failure, and stroke.^1,3–5^ AF pathogenesis is complex, multifactorial, and poorly understood. Abnormalities in cellular electrophysiology, impaired Ca^2+^ homeostasis, mitochondrial dysfunction, inflammation, and genomic instability with dysfunctional DNA repair mechanisms have all been suggested to be involved in AF pathophysiology.^2,6–11^ Systemic processes such as cardiorenal syndromes may contribute to AF initiation and progression.^12,13^ Regardless of severity, chronic kidney diseases significantly increase the risk of incident AF.^14^

Mutations in Polycystin-1 (PKD1, hereinafter PC1) are the primary culprit of autosomal dominant polycystic kidney disease (ADPKD) cases, a disease characterized by the formation of fluid-filled cysts that expand and degenerate over time, leading to renal failure. A higher-than-expected frequency of *PKD1* mutations in healthy sequenced populations has been recently described^15^, and a large genomic study revealed that *PKD1* mutations have a 47% penetrance for chronic kidney disease.^16^ This evidence suggests that PC1 mutations are more common than originally thought. Although primarily classified as a renal pathology, cardiovascular disease is the leading cause of morbidity and mortality among ADPKD patients. These patients have increased rates of hypertension, vascular alterations, left ventricular hypertrophy, diastolic dysfunction, reduced left ventricular strain, dilated cardiomyopathy, AF, and other arrhythmias.^3,17–21^ The current dogma is that cardiovascular alterations in ADPKD arise due to renal complications. However, new evidence indicates subclinical cardiovascular alterations in ADPKD patients with normal renal function.^22–26^ Indeed, a recent study from the US-National Inpatient Sample database, including 71,531 patients, revealed that a higher risk of AF and atrial flutter was observed among ADPKD in the younger age group (18-39 years of age) compared to those with CKD and without ADPKD (odds ratio of 5.71).^27^ Given that ADPKD increases the risk of AF ^21,23,27–29^, we investigated whether mutations in PC1 contribute to increased AF inducibility.

PC1 is not only implicated in ADPKD but is also ubiquitously expressed across multiple tissues, including the heart, showing its broader biological significance. Evidence points out that PC1 plays an important role in cardiomyocytes. We and others have shown that PC1 is essential in ventricular cardiomyocyte excitation-contraction (EC) coupling, hypertrophic response to pressure overload, mitochondrial function, and cardiac fibrosis.^30–33^ However, no studies have directly elucidated the role of PC1 in the atria. Moreover, little is known about the effects of PC1 on transcription regulation in the heart, as its C-terminal tail regulates gene expression in various cell types.^34–36^ Here, we sought to investigate the role of PC1 in atrial cardiomyocyte function and its impact on AF susceptibility, employing multiple mouse and cellular models. We hypothesized that PC1 directly regulates atrial cardiomyocyte function and protects against AF. Unexpectedly, we found that PC1 regulates genes involved in DNA repair. PC1 deficiency increases susceptibility to DNA damage and arrhythmias. This is supported by recent evidence that proposes DNA damage as a novel pathway contributing to AF progression.^6,8,11,37^ Our work suggests that PC1 protects against DNA damage and AF susceptibility.

## Methods

Additional detailed methods are presented in supplementary material online.

### Animal models

All experiments were performed following institutional guidelines and approval by the Institutional Animal Care and Use Committee of the Houston Methodist Research Institute. Cardiomyocyte-specific PC1 KO (cKO) was generated as outlined in supplementary methods. *Pkd1*^RC/RC^ mouse line was obtained from Dr. Peter Harris at Mayo Clinic, MN, USA. Both male and female mice were used for experiments at 2-4 months of age. The study was not specifically designed to detect or quantify gender-based differences.

### Intracardiac Electrophysiology

Programmed intracardiac electrical stimulation was performed using an octopolar catheter in mice, as previously described.^38–41^ AF susceptibility was assessed using three rounds of 2-second bursts for each cycle with an initial cycle length of 40 ms and decrementing cycle length by 2 ms, for a final cycle length of 10 ms. Thirty seconds of recovery were allowed between each burst pacing. *In vivo* AF rates were identified as rapid, irregular signals in the atrial electrode, the absence of P waves, and increased R-R variability. No ventricular arrhythmias were observed before or after programmed electrical stimulation.

### Optical Mapping

A 530 nm LED source and MiCAM Ultima acquisition system were used to capture images from atrial left appendages loaded with RH-237 and Rhod-2 AM at 500 fps. Monolayers were mapped using Rhod-2 AM at 200 fps.

### Cellular Models

Human episomal iPSC cell line (A18945, hPSCreg Name TMOi001-A) was purchased from ThermoFisher and differentiated into atrial cardiomyocytes (hiPSC-aCMs) using retinoic acid as previously described.^42^ Following differentiation, cells were passaged onto new Matrigel (Corning) coated plates and maintained in culture media RPMI supplemented with B27. Experiments were performed after day 35. HL-1 cells were obtained from Sigma Aldrich (SCC065) and cultured in Claycomb media as previously described.^43,44^

### Multielectrode Array (MEA)

Signals were recorded using an 8×8 grid of carbon nanofiber electrodes (100 μm x-y spacing) using MED64 Presto (Alpha Med Scientific, Osaka, Japan).

### Assessment of action potentials and intracellular Ca^2+^

A modified IonOptix system was used to detect FluoVolt, Fura-2AM or Fluo-4AM signals during pacing at 2 Hz using field electrical stimulation.

### RNA Sequencing

Fully unbiased bulk RNA sequencing and preliminary bioinformatic analysis were performed by GeneWiz (NJ, USA). RNA counts were then used to identify differentially regulated pathways using overrepresentation analysis (ORA) and Gene Set Enrichment Analysis (GSEA) using clusterProfiler v4.0 ^45^, Ingenuity Pathway Analysis (IPA Qiagen) and pyDESeq2/GSEApy/decoupler 2.

### Human Samples

Left atrial appendages were obtained from sinus rhythm and persistent AF patients undergoing open-heart surgery as previously described.^46,47^ AF characteristics were determined based on clinical information (Table S6). Experimental protocols were approved by the ethics committee of the University Medical Center Göttingen (No. 4/11/18) and were performed in accordance with the Declaration of Helsinki. Written and informed consent was given by each patient before surgery.

### Statistical Analysis

All statistics were performed using GraphPad Prism Software and R Studio 2023.03.0 Build 386/R version 4.2.3. Categorical data were analyzed using Fisher’s exact test. Normality was tested using the Shapiro-Wilk test. Quantitative variables were compared using two-tailed t-test or Mann-Whitney test as appropriate. For the comparison of more than two groups, a one-way or two-way ANOVA was performed with Dunnett’s correction for multiple comparisons. A nested t-test was also applied for experiments with multiple cells/batches. A *P* < 0.05 was considered statistically significant. Comprehensive details of statistical analyses for each figure are provided in Table S1.

### Data Availability

The data supporting this study’s findings are available from the corresponding author upon request. RNA-seq raw counts will be uploaded to NIH GEO before publication. Supplemental Material presents an expanded version of the Methods section and Supplemental Figures, Tables, and Videos.

## Results

### Loss of PC1 function increases *in vivo* AF susceptibility

We examined AF inducibility in knock-in *Pkd1*^RC/RC^ mice – a well-validated mouse model that recapitulates human renal ADPKD progression and harbors p.R3277C mutation, herein after RC.^48–50^ The RC mutation is a temperature-sensitive folding mutant with impaired endoplasmic reticulum to trans-Golgi trafficking, which affects PC1 maturation (glycosylation pattern) and protein sorting.^50^ RC mice renal disease progression recapitulates human ADPKD: formation of renal fluid-filled cysts occurs at the embryonic stage with a progressive postnatal increase in kidney volume (significantly different at 3 months), and the onset of renal failure at 9 months of age.^48,50^

We performed experiments between 2-4 months of age to mitigate the consequences of renal disease impinging on atrial function. Our data shows that young adult RC mice exhibit a higher incidence of AF after atrial burst pacing, compared to WT mice (Fig. 1A-D, S1, and S7, refer to Table S1 for all statistical analyses). No major cardiac changes were observed in the morphometric analysis (heart weight and atrial size), and both atrial and ventricular histology were normal by hematoxylin and eosin (H&E) and Masson’s trichrome staining in WT and RC mice (Fig. S2-5). Additionally, normal cardiac function was assessed by echocardiography in RC mice (fractional shortening and diastolic function, Fig. S7). Although we observed a slight reduction in fractional shortening in RC mice, histology and ventricular dimensions obtained by echo and histology suggest no evidence of cardiac adverse remodeling. Similarly, we did not observe changes in diastolic function measured by pulse wave and tissue Doppler (E/e’). This evidence suggests that increased susceptibility to AF in RC mice does not arise from ventricular remodeling or increased filling pressure. We observed slight increases in kidney weight corrected by body weight and evidence of renal cysts and fibrosis, as expected and reported in RC mice^50^ before the onset of renal failure (Figure S2-3, and S6). Together, our findings show that one single amino acid substitution in the highly characterized PC1 RC mutation increases AF vulnerability before the onset of renal failure.

**Figure 1:**
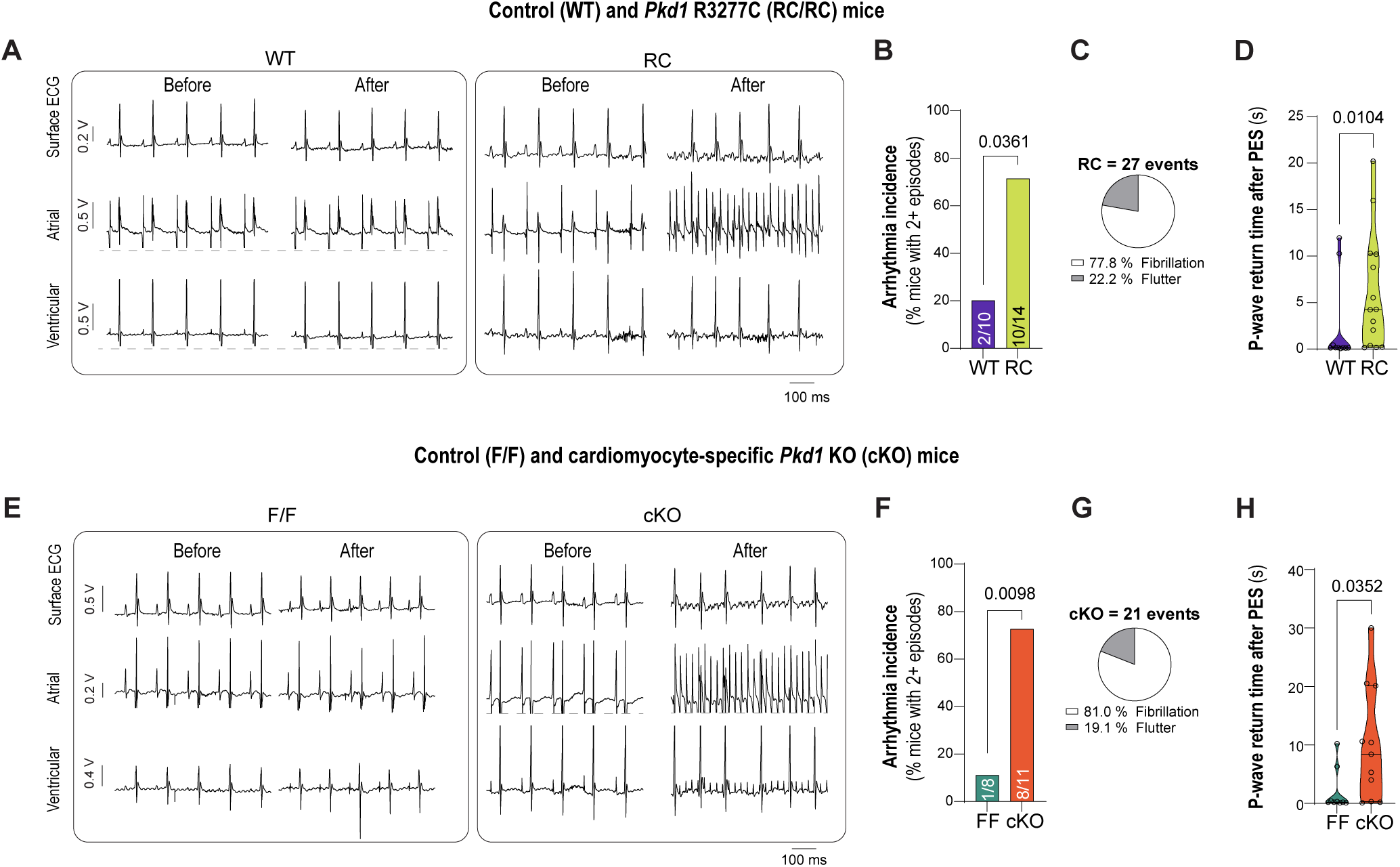
PC1 deficiency increases vulnerability to atrial fibrillation. Intracardiac electrophysiology and programmed electrical stimulation (PES) were performed in young (2-4-month-old) mice. A total of 3 challenges were performed with a resting interval of 30 s. *Top panel* (**A-D**): *WT control* vs. *Pkd1* R3277C (ADPKD preclinical model). *Bottom panel* (**E-H**): *Pkd1*^F/F^ (WT control with floxed alleles) vs. *Pkd1*^F/F^ *Myh6*-Cre (cardiomyocyte-specific PC1 KO) mice. **A** and **E**, representative recordings showing surface electrocardiograms and intracardiac atrial and ventricular leads obtained with an octopolar catheter advanced through the jugular vein. **B** and **F**, higher arrhythmia incidences were observed in PC1-deficient mice. The number of mice with >2 arrhythmic events over the total mice is indicated at the bottom of each bar. **C** and **G**, in PC1-deficient mice, we observed numerous arrhythmia events. All of these events were atrial arrhythmias, with the majority being atrial fibrillation: disorganized high-frequency atrial signals and the disappearance of P-wave in surface ECG. A smaller number of events were atrial flutter (multiple P-waves with regular atrial signals). **D** and **H**, Duration of the longest arrhythmia measured as P-wave return time after programmed electrical stimulation (in mice without arrhythmias this period typically takes <0.5 second, as the baseline ECG takes a few seconds to stabilize and show the first clearly visible QRS complex after program electrical stimulation). Data are presented as violin plots, including individual data points, median and upper and lower quartiles. Exact P-values are shown on each graph, ns; non-significant difference. <u>Only one</u> <u>mouse was excluded for excessive bleeding during the procedure</u>. Data were collected and analyzed blinded for genotype. Additional information regarding N number and statistical analysis can be found in Table S1a.

To exclude extra-cardiomyocyte effects causing AF, we then studied AF vulnerability in cardiomyocyte-specific PC1 KO (*Pkd1*^F/F^; *Myh6*–*Cre*; cKO), a model that does not directly impair kidney function, and control (*Pkd1*^F/F^; F/F) mice. Similar to our findings in the RC model, young cKO mice (2-4 months old, of either sex) showed increased AF susceptibility after atrial burst pacing compared to F/F mice (Figure 1E-H, S1, and S7). These observations show that PC1 protects against AF in mouse models, likely due to a cardiomyocyte-specific mechanism. Thus, we aimed to elucidate further the direct contribution of PC1 in atrial cardiomyocytes and arrhythmias using a multi-strategic approach.

### Increased local reentry in the left atrial appendage of RC mice

Voltage and Ca^2+^ optical mapping were used to study atrial arrhythmogenesis in superfused left atrial appendages (LAA) from WT and RC mice. Tests were conducted sequentially at frequencies between 6 and 12 Hz, with each frequency maintained for 30 seconds to reach a steady state before recording for 2 seconds at 500 fps. In 3 out of 4 RC mice, unidirectional conduction block and local reentry circuits were observed, whereas WT mice exhibited normal responses.

Fig. 2A and Supplemental Videos 1 and 2 show side-by-side comparison and quantification of Ca^2+^ optical mapping data sequentially obtained between 6-12 Hz in superfused LAA from WT and RC mice. Homogeneous impulse propagation was observed at all tested frequencies in WT tissue (Fig. 2A-D), and frequency analysis showed single frequency domains in all tissues, matching the pacing frequency. On the contrary, RC tissue showed dramatic abnormalities starting at 10 Hz, demonstrated by frequency response breakdown and multiple frequency domains (Fig 2A, E-G and Supplemental Video 2). Local reentry was observed in RC mice, and it was triggered from specific areas with delayed conduction that facilitated local reentry during interbeat, which then merged with subsequent pacing signals (Fig 2G). Among the RC mice demonstrating abnormal responses, 2/3 exhibited similar patterns of local reentry and conduction block. Interestingly, the third tissue displayed these aberrations at lower frequencies (9-10 Hz), which led to conduction blocks in the affected areas when exposed to higher frequencies.

**Figure 2:**
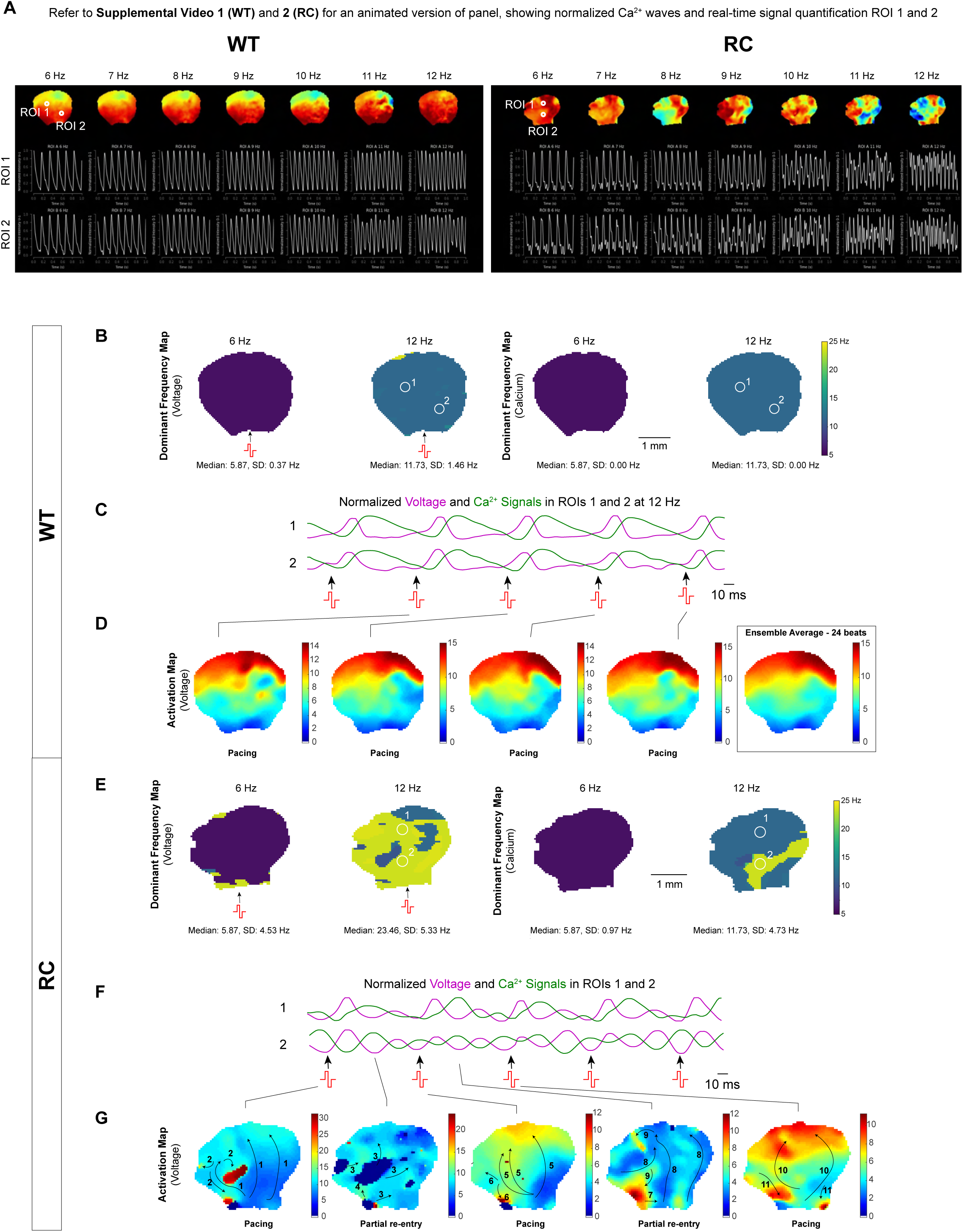
Influence of PC1 R3277C (RC) mutation on atrial arrhythmias. **A.** Ca^2+^ optical mapping of superfused left atrial appendages (LAA) obtained from RC mice show increased susceptibility to tachyarrhythmia compared to WT counterparts. Animated version of these figures are provided in Supplemental Video 1 and 2 for WT and RC, respectively (refer to Supplemental Video Legend for more information). The tissues were sequentilly paced and tested at frequencies between 6-12 Hz using a pair of platinum wires located at the bottom of the tissue. The normalized signal (ranging from 0-1) is shown with blue to red scale. Real-time quantifications from two regions of interest (ROI 1 and 2; highlighted in top panels) are displayed in top and bottom plots. **B**. Voltage and Ca^2+^ frequency maps, with dominant frequencies calculated with Fourier Fast Transform show normal responses in WT tissue. **C**. Signal comparison in WT tissue: Normalized voltage (purple) and Ca^2+^ (green) signals in marked ROIs above for control show a regular rate with voltage signal preceding Ca^2+^. Pulses marked by biphasic red waveform and arrowheads. **D**. Voltage activation maps for WT atria show even activation of the whole mouse heart. Color scale bar units = ms. **E**. Voltage and Ca^2+^ maps showing dominant frequency calculated with Fourier Fast Transform in RC tissue. A disordered and local increase in dominant frequency across the atria was observed starting at 10 Hz and shown at 12 Hz point stimulation (for voltage and Ca^2+^ signals). **F**. Similar to Panel **C**, but for RC tissue, demonstrating altered frequency and timing of signals. **G**. Voltage activation maps highlight areas of conduction delay and partial reentry in RC mouse. Color scale bar units = ms.

To investigate the underlying mechanisms, we examined Connexin 40/43 expression in LAA tissue slides obtained from WT and RC mice. CellProfiler software was used to measure clustering and fluorescence intensity in an unbiased manner (Figure S8). Our analysis revealed no significant differences in connexin clustering or expression levels. Together with previous data indicating the absence of fibrosis, these findings suggest that the observed AF-like episodes are not attributable to connexin remodeling, fibrotic changes or ventricular dysfunction, suggesting alternative mechanisms driving these alterations.

### PC1-deficiency increases *in vitro* arrhythmias

Confluent hiPSC-aCMs monolayers were plated on multi-electrode array (MEA) plates and allowed to synchronize. We then transiently knocked down PC1 expression by ∼75% using siRNA and compared our data with monolayers transfected with control siRNA (Figure 3). PC1-deficient hiPSC-aCMs showed increased nonpropagating arrhythmic events (measured as ectopic beats, regional conduction slowing, and local conduction block) compared to control (Figure 3C-F). These local arrhythmic events were limited to a few electrodes and did not change average conduction velocity or beating frequency; however, we observed a significant reduction in field potential duration (FPD) (Figure 3G-J). Optical mapping studies also revealed that PC1 silencing increased spiral wave activity or rotors under baseline conditions, aggravated by tachypacing (see Figure 5 and next sections). These findings suggest that PC1 deficiency leads to *in vitro* arrhythmias in atrial cardiomyocytes.

**Figure 3:**
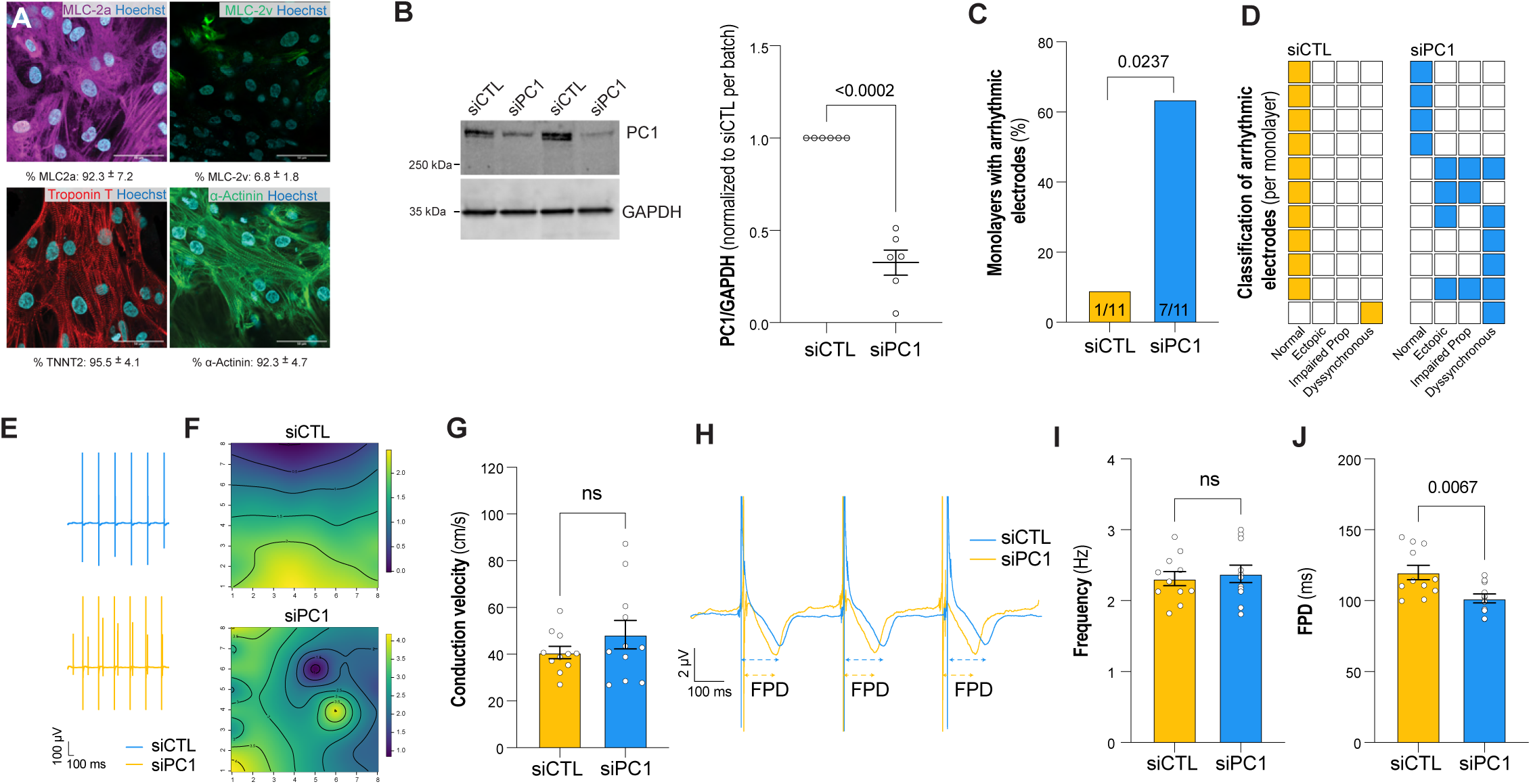
Increased *in vitro* arrhythmias in PC1-deficient atrial cardiomyocytes. **A.** We stained for atrial and ventricular myosin light chains (MLC2a and MLC2v, respectively) and common cardiac markers (Troponin T and α-actinin). The percentage of positive cells is provided under each representative figure. Cells were used >35 days after differentiation. **B.** hiPSC-aCMS were transfected using CTL or PC1 siRNA (siCTL and siPC1). Knockdown efficiency was, on average, between 70-80% (refer to Figure 6I) to knock down its protein levels by 70%. **C**. hiPSC-aCM were plated in MEA plates with 8×8 carbon nanotube electrodes, and the incidence of arrhythmias was measured after transfection with siCTL (**blue**) and siPC1 (**orange**) at 48h. PC1-deficient monolayers exhibited increased arrhythmic events compared to control (N=11 monolayers per condition). **D.** Classification of arrhythmic events based on ectopic activity, impaired propagation (>10 ms), and dyssynchronous propagation/conduction block electrodes. **E.** Representative traces showing ectopic activity in PC1-deficient cells. **F.** Spline surface plots were calculated to visualize electrical signal propagation in the 8×8 grid of electrodes. Representative Figure showing electrodes with dyssynchronous propagation/conduction block after PC1 ablation. **G.** No changes in conduction velocity were observed between monolayers treated with CTL and PC1 siRNA. **H.** Representative field potentials showing similar (**I**) beating frequency (**J**) but shorter field potential duration (FPD) in PC1-deficient cells.

### PC1 regulates the expression of DNA repair genes

PC1 has previously been shown to be a transcriptional regulator of multiple cellular pathways, primarily through its C-terminal tail cleavage and subsequent transcriptional regulation.^34–36^ We hypothesized that PC1 regulates crucial pathways in atrial cardiomyocytes, potentially influencing AF susceptibility. To test this hypothesis, we analyzed the transcriptional profiles of hiPSC-aCM transfected with control and PC1 siRNA (siCTL and siPC1, respectively) at 48h obtained from five independent differentiation batches assessed in duplicates using RNA-seq.

We identified 103 downregulated and 2 upregulated genes in siPC1 compared to siCTL atrial cardiomyocytes (adjusted p-value or FDR <0.05 and absolute value of log_2_FC > 1, Figure 4A and Table S1). For gene ontology analysis, we included genes with FDR < 0.05 and an absolute value of log_2_FC > 0.5 (196 downregulated and 28 upregulated). Gene Ontology Biological Process (GO: BP) analysis revealed that siPC1 decreases the expression of multiple genes associated with DNA repair, mitosis checkpoints, and cytoskeleton/chromosome regulation (Figure 4B and Table S2).

**Figure 4:**
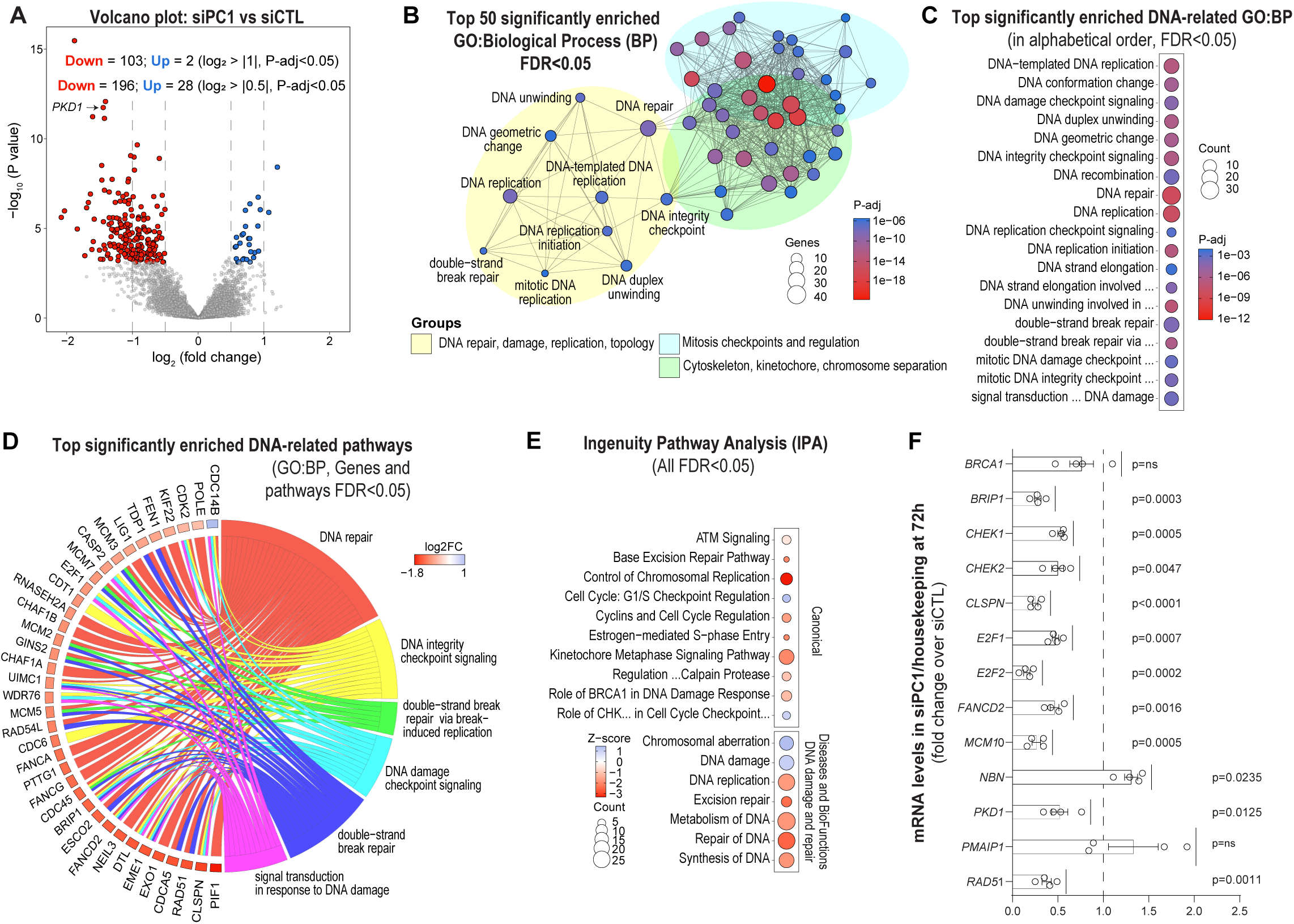
PC1 regulates the expression of DNA repair genes. We performed unbiased transcriptomics profiling using Genewiz RNA-seq service (see Methods). **A**. Volcano plot showing differentially expressed genes (DEGs). Refer to Table S2 for more details. **B**. Genes with FDR < 0.05 and absolute value of log_2_FC> 0.5 were used for overrepresentation analysis using clusterProfiler v.4.0. Top 50 enriched Gene Ontology (GO) - Biological Process (BP) clustered in 3 main groups, including DNA repair pathways. Circle size represents the number of DEGs enriched for each pathway (count) and statistical significance is represented with a red-to-blue color scale (p-adjusted or FDR). Data in bubble plots are statistically significant (FDR < 0.05) and presented alphabetically. **C-D**. Top significantly enriched DNA-related GO-BP and Circos chord plot showing the relationship between DNA-related GO-BP terms and DEG. Red-white-blue scale shows log_2_FC for genes with FDR < 0.05. **E.** We then used ingenuity pathways analysis (IPA, Qiagen) to predict pathway activation or inhibition (Z-score is provided for significant pathways; FDR<0.05). Sample processing and statistical analysis (DESeq2) were fully unbiased and performed by GeneWiz. FDR was calculated using the Benjamini-Hochberg test in DESeq2, clusterProfiler and IPA. Refer to Table S1 and S3a for more details. **F**. Verification of RNA sequencing data was partially assessed using qPCR. Data were normalized using 4 housekeeping genes (GAPDH, HPRT1, PGK1, and TBP). Refer to Table S1 for additional details about the statistical analysis.

Recent studies suggest a connection between DNA damage and AF pathogenesis. Transient tachypacing leading to AF episodes increases DNA damage by reducing NAD^+^ levels, impairing mitochondrial function, and increasing ROS levels.^11^ In addition, a stepwise increase in DNA damage has been observed in paroxysmal and persistent AF.^37^ We discovered that PC1 regulates essential genes involved in DNA repair, which were downregulated in siPC1 compared to siCTL (Figure 4C-F, Table S2-3). Furthermore, pathway activation predictive analysis using Ingenuity Pathway Analysis (IPA) software prediction reveals a down-regulation of DNA repair pathways and increased DNA damage pathways (Figure 4E). In simpler terms, when PC1 is silenced, the expression of multiple genes associated with DNA repair decreases, which results in increased susceptibility to DNA damage under stress conditions. This evidence suggest that PC1 protects atrial cardiomyocytes against DNA damage (see below).

To corroborate our results, we measured 1) RNA levels using qPCR and observed a reduction in the expression of key regulators of DNA repair pathways (Figure 4F), and 2) protein levels using western blot and found reduced protein levels of CHK1, TOP2a, CEP55, MCM5 and RAD51 (Figure S9A), confirming that PC1 deficiency decreases the expression of key regulators involved in DNA damage/repair responses. A similar result was observed in cKO total atria lysates, showing significant downregulation of TOP2a and nonsignificant downregulation of Chk1 (P=0.061) and Rad51 (P=0.126) compared to F/F (Figure S9B).

### PC1 protects against atrial arrhythmias through the regulation of DNA repair pathways

Recent studies have shown a link between DNA damage and AF, highlighting the role of excessive poly ADP-ribose polymerase (PARP1) activity.^11^ This excessive PARylation contributes to the depletion of nicotinamide adenine dinucleotide (NAD^+^), resulting in significant metabolic disturbances and compromised EC coupling.^8,11^ We hypothesized that loss of PC1 increases susceptibility to DNA damage, enhancing susceptibility to atrial arrhythmias. To demonstrate mechanistic connections between PC1 levels, DNA damage, and arrhythmias, we used tachypacing as an *in vitro* AF model.^11^

Monolayers of hiPSC-CM were treated with siCTL and siPC1 and were then tachypaced using field electrical stimulation (see Methods). DNA damage was assessed by γH2AX staining. At baseline, PC1-deficient hiPSC-aCM showed a slight increase in DNA damage, but this difference was not statistically significant. This damage was further exacerbated after tachypacing (Figure 5A) and more so in the PC1 siRNA-treated monolayers. This evidence suggests impaired PC1 increases susceptibility to DNA damage under stress conditions, e.g., tachypacing. In RC atrial tissue sections, we found a significant increase in the baseline γH2AX fluorescence speckle intensity and size compared to the control and clustering of DNA-damaged cardiomyocytes within the atria (Figure 5B and S8B).

**Figure 5:**
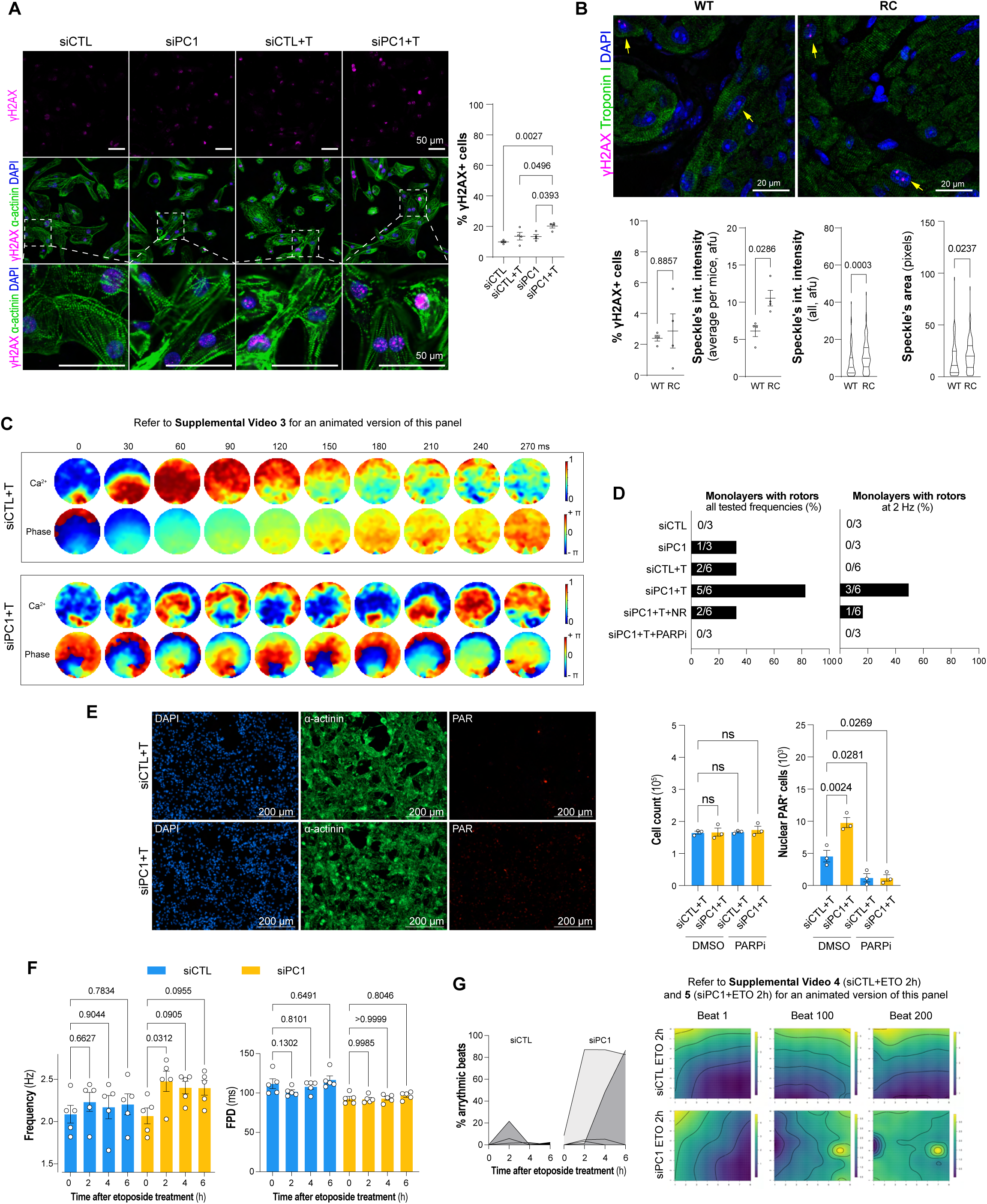
PC1 protects against DNA damage and tachypacing-induced arrhythmias. **A.** Monolayers were transfected 48 h prior the experiment with CTL or PC1 siRNA (siCTL or siPC1, respectively). Tachypacing was performed as described in Supplemental Methods. **A.** Increased levels of γH2AX levels, indicating DNA damage, were observed under basal conditions and significantly increased in PC1-deficient cells as early as 4 h post-tachypacing. **B.** Atrial tissue from RC mice exhibits greater DNA damage than control, as shown by increased speckle integrated intensity and area measurements. **C**. Optical mapping of Ca^2+^ signals revealed spontaneous spiral-wave reentry in PC1-deficient cardiomyocytes, which was further aggravated by tachypacing (Supplemental Video 3, refer to Supplemental Video Legend for more information). We assessed arrhythmias in monolayers by subjecting them to various stimulation frequencies (2-6 Hz point stimulation). Our main focus was on the presence of rotors, which are spiral waves indicating arrhythmia. These rotors were observed variably and were not visible at all frequencies, highlighting the unpredictable nature of arrhythmias. **D**. We classified monolayers as arrhythmic if a rotor was detected at any frequency. PC1 deficiency increased rotor susceptibility after tachypacing, even at lower frequency stimulation. This effect was partially rescued by NR, with no rotor activity with PARPi. **E**. Tachypaced monolayers were fixed and stained for α-actinin, Poly-ADP Ribose (PAR), and counterstained with DAPI to detect nuclei. A total of 15×15 stitched images obtained with a 10x objective were obtained with Cytation 1. Representative image showing increased nuclear PAR staining in PC1-deficient cardiomyocytes. The total number and PAR^+^ cells per monolayer was measured in 15×15 fields using a high-throughput microscope (Cytation 1) and Gen 5 software (a threshold was used by measuring background/signal intensity to identify positive cells). PC1 deficiency increased the number of PAR+ cells, blunted by PARP1 inhibition with Veliparib (ABT-888, 20 μM). **F**. Cells were plated in MEA plates and then exposed to 10 μM etoposide. Electrophysiological measurements were obtained before and after 2, 4, and 6 h post-treatment. Frequency and field potential duration (FPD) are shown. **G.** PC1-deficient monolayers exhibited an increase in arrhythmic activity at 2 h, which remained elevated throughout 6 h, whereas CTL had a slight transient increase in arrhythmic behavior (Supplemental Video 4 and 5, refer to Supplemental Video Legend for more information). Data are presented as the mean value ± SEM, and the individual data points are also shown (with the exception of categorical data). Refer to Table S1b for additional information.

We then performed Ca^2+^ optical mapping in monolayers treated with siCTL and siPC1 at baseline and after tachypacing. We evaluated responses to frequency (2-6 Hz) through point stimulation, and we found increased arrhythmic rates in PC1-deficient monolayers compared to control, even at lower tested stimulation frequencies (Figure 5C-D). Interestingly, siPC1 increased spiral wave reentry with phase singularities, as depicted in Figure 5C and Supplemental Video 3. To determine whether the consequences of increased DNA damage in PC1 deficient monolayer trigger arrhythmias, we use nicotinamide riboside (NR) supplementation and PARP1 inhibition (ABT-888 or Veliparib). Remarkably, treatment with these molecules corrected arrhythmic rates in PC1-deficient atrial cardiomyocytes (Figure 5D). Furthermore, tachypacing led to a 2-fold increase in the abundance of PAR-positive cell abundance in siPC1 compared to siCTL cells, without affecting cell viability. This was detected using a fluorescence intensity threshold to identify PAR signal above background noise, which colocalized with nuclear masks (DAPI staining; average of 15×15 stitched images obtained at 10X magnification). Notably, this increase was significantly mitigated by PARP1 inhibition (Figure 5).

To validate our findings, we performed additional experiments: 1) We directly tested whether DNA damage increases vulnerability to arrhythmias in siPC1 using etoposide treatment (10 μM). Etoposide induces DNA breaks by inhibition of topoisomerase II (most abundantly expressed in cardiomyocytes is TOP2b, which was not affected by siPC1). PC1 deficiency increased susceptibility to etoposide-induced arrhythmias as measured by increased beating frequency and the percentage of arrhythmic beats (Figure 5F-G and Supplemental Videos 4 and 5). 2) In HL-1 cells, etoposide further increased DNA damage and arrhythmia rates in PC1-deficient cells (Figure S10). Furthermore, we performed γH2AX clearance assays to reveal that loss of PC1 impairs DNA repair in atrial cardiomyocytes (Figure S10H). Taken together, our findings suggest that loss of PC1 increases vulnerability to DNA damage through impaired DNA repair.

### PC1 regulates DNA repair genes through its C-terminus tail

Cleavage of PC1 is known to regulate the activity of multiple transcription factors likely via nuclear translocation of its C-terminus tail (PC1-CT).^35,51,52^ Interestingly, in adult mouse atrial cardiomyocytes and hiPSC-aCM, the endogenous PC1-CT localized with both striated and nuclear pattern (S11A-B). Staining specificity was demonstrated in adult mouse atrial cardiomyocytes from cKO mice and hiPSC-aCM treated with siPC1 (Figure S1D and S11E). Since protein sequences for mouse and human PC1-CT differ, these observations were made with highly validated antibodies developed by PKD Cores at the University of Maryland (Kerafast #EJH002) and Kansas (161F^53^, see acknowledgments), respectively. A similar distribution pattern was observed when expressing a fused mCherry-PC1-CT-V5tag using adenovirus, which expresses cytosolic PC1 C-terminus tail. Live-cell mCherry fluorescence and co-staining of mCherry and V5 in fixed cells showed striated and nuclear localization (Figure S11C-D). The association between PC1 and the actomyosin complex and nuclear localization of PC1-CT has been described in other cell types.^52,54^ Interestingly, overexpression of PC1-CT was able to recapitulate the striated and nuclear distribution of the endogenous protein in hiPSC-aCM. Although our studies uncover PC1-CT localization, multiple cleavage sites have been reported in PC1 within transmembrane domains and the short cytoplasmic tail (reviewed in ^52^). More studies are required to determine the exact cleavage(s) releasing PC1-CT in atrial cardiomyocytes.

To gain a better understanding of how PC1 regulates DNA damage, we sequenced RNA from hiPSC-aCM that overexpress the PC1-CT using adenovirus, with LacZ as a control. PC1-CT modified the expression of 105 genes (77 down and 28 up-regulated, FDR <0.05 and an absolute value of log_2_FC > 1; Figure 6A, Table S4). Co-expressed Zsgreen1 under a second CMV promoter in adenoviral constructs demonstrates a 100% infection efficiency (Figure 6B). We noted that multiple genes followed opposite trends after PC1 knockdown or overexpression. To further assess PC1 effects in hiPSC-aCM, we independently analyzed our two RNA-seq datasets (#1 siPC1 vs. siCTL presented in Figure 4A; #2 PC1-CT vs LacZ presented in Figure 6A) using Gene Set Enrichment Analysis (GSEA, Figure 6C, Table S5a and S5b). We first created two custom gene sets obtained from the leading edges of each dataset: downregulated genes in dataset #1 ("PC1 KD Down"; FDR <0.05 and log_2_FC <-0.5; 196 genes) and upregulated genes in dataset #2 ("PC1 CT Up"; FDR <0.05 and log_2_FC >0.5; 335 genes).

**Figure 6:**
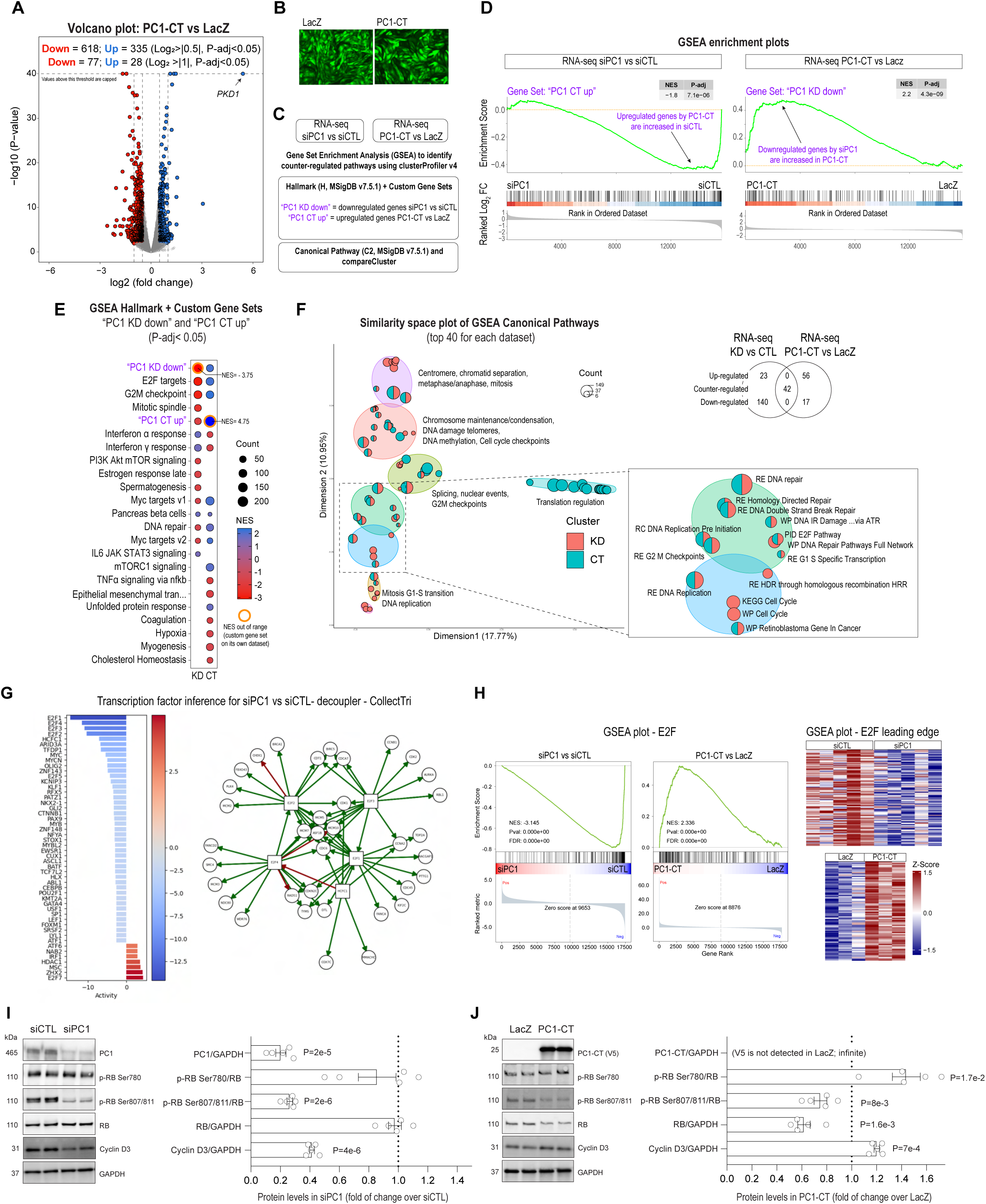
PC1 regulates the expression of DNA damage repair genes via its C-terminal tail. We used adenovirus to express PC1 C-terminus (PC1-CT or CT). Total RNA was extracted, and an unbiased transcriptomics analysis was conducted by GeneWiz. **A**. Volcano plot showing differentially expressed genes (DEGs). Refer to Table S4 for more details. **B**. Zsgreen1 fluorescence showing 100% infection efficiency in cells treated with LacZ and PC1-CT adenoviruses. **C**. We evaluated the effects of PC1 manipulation in hiPSC-aCM by comparing our RNA-seq databases using GSEA as described in the figure. **D**. Using the compareCluster function (clusterProfiler v4.0), we performed GSEA using our custom gene sets in combination with MSigDB – Hallmark gene sets (v.7.5.1) in both RNA-seq databases. These gene sets exhibit counter-regulation when analyzed in the opposite RNA-seq database. **E.** Furthermore, GSEA revealed multiple counter-regulated pathways, including E2F targets, G2M checkpoints, and DNA repair (Table S5a). **F**. We then expanded our GSEA analysis to MSigDB – C2 canonical pathways gene sets (v.7.5.1). shows various clusters of counter-regulated pathways regulated by PC1. Indeed, multiple pathways associated with DNA damage repair were downregulated in PC1-deficient cells and upregulated after PC1-CT overexpression, as shown in the similarity space plot (Table S5b). **G**. Transcription factor inference for siPC1 vs siCTL using decoupler 2 and CollecTRI database showing E2F inhibition and network of downregulated genes. **H**. GSEA plots obtained using GSEApy using stat for gene ranking show similar results to the ones obtained with clusterProfiler. E2F downregulation is observed in siPC1, whereas it is enhanced by PC1-CT. Representative western blot and quantification showing RB phosphorylation and total levels in siCTL and siPC1 and LacZ vs PC1-CT (**I** and **J**, respectively). Additional information regarding the statistical analysis can be found in Table S1a.

Using compareCluster (clusterProfiler v4.0 ^45^), we performed GSEA with our custom gene sets in combination with MSigDB – Hallmark gene sets v.7.5.1, in both RNA-seq datasets. Remarkably, these gene sets were significantly counter-regulated when assessed in the other dataset. In other words, genes downregulated by PC1 deficiency were upregulated by PC1-CT overexpression and vice versa (Figure 6D). Furthermore, GSEA revealed multiple counter-regulated pathways, including E2F, G2M checkpoints, and DNA repair – all pathways related to DNA damage responses (Figure 6E).

Expanding our GSEA to include MSigDB – C2 canonical pathways gene sets (Figure 6F), we observed significant counter-regulated pathways in clusters related to DNA repair. Specifically, PC1 deficiency led to a downregulation of genes crucial for DNA repair pathways, whereas overexpression of PC1-CT showed an upregulation of these genes. This suggests that while PC1 does not directly induce DNA damage pathways, it plays a critical regulatory role in modulating the expression of key genes involved in responding to DNA damage and facilitating DNA repair processes.

GSEA Halmark and decoupleR 2/CollectTRI enrichment analysis revealed insights into the regulatory transcriptional networks regulated by PC1 (Figure 6G; also observed with TRRUST; data not shown). Transcription factor inference suggests the reduced E2F activity in PC1-deficient cells, which is associated with the downregulation of multiple genes (Figure 6G, right panel, green arrows) in siPC1 vs siCTL. These findings are corroborated by GSEA analyses showing E2F geneset downregulation in siPC1 vs siCTL and upregulation in PC1-CT vs LacZ (Figure 6H). This pattern of opposing regulation suggests that PC1-dependent regulation of E2F may be involved in impaired DNA repair in atrial cardiomyocytes (Figure 6E and 6G-J). Specifically, the Retinoblastoma (RB)/E2F pathway is critical for DNA damage responses, as it regulates multiple genes required for repair^55,56^ and also directly facilitates DNA repair at damage sites.^57–59^ Decreased E2F activity is likely due to decreased RB phosphorylation at Ser805/811 (Figure 6I) in siPC1 compared to siCTL. In its hypophosphorylated form, RB binds to E2F, preventing E2F from activating gene targets. On the contrary, increased E2F transcriptional activity in PC1-CT vs LacZ, was likely caused by a significant decrease in RB total protein and increases in Ser780 phosphorylation (Figure 6E, H and J). Although a slight reduction in Ser805/811 was observed, nearly 50% of RB’s total mass was decreased, suggesting E2F activation.

Although the exact mechanism by which PC1 regulates E2F is not known, our data shows a differential regulation of cyclin D3 (Figure 6I-J), which plays a biological role in adult non-dividing tissues to maintain a terminally differentiated phenotype.^60,61^ Cyclin D-types activate Cdk4/6 to promote RB phosphorylation.^60^

Future studies are needed to elucidate this molecular mechanism further as the RB phosphorylation code is complex and integrates E2F association with multiple partners regulating the diversity of RB activity in DNA repair, proliferation, and metabolism.^62^ Given that E2F1, a member of the E2F family, downregulation increases AF susceptibility,^63^ it is possible that PC1-dependent regulation of E2F1 maintains DNA integrity in atrial cardiomyocytes to protect against AF. Interestingly, recent studies show adverse cardiovascular events (CVAEs) associated with cyclin-dependent kinase (Cdk) 4/6 inhibitors in patients with metastatic breast cancer. CVAEs were slightly higher in patients who received Cdk4/6 inhibitors compared with anthracyclines, with a higher death rate associated with the development of atrial fibrillation/flutter or cardiomyopathy/heart failure in the Cdk4/6 group.^64^ These findings highlight the relevance of the studied pathway.

### Alternative mechanisms regulating arrhythmia vulnerability in PC1 deficient atrial cardiomyocytes

We measured action potentials using the potentiometric dye FluoVolt during steady-state pacing at 2 Hz using a modified IonOptix imaging system. siPC1 significantly shortened APD at 50 and 80% repolarization compared to siCTL (Figure 7A). The observed shortening of FPD and APD in hiPSC-aCM following PC1 silencing (Figure 3 and 7A) could likely be attributed to increased Kv4.3 and Kv1.5 currents as we previously showed that the human version of PC1 negatively regulates macroscopic currents of human Kv4.3 and Kv1.5, which are abundantly expressed in human atria.^30^

**Figure 7:**
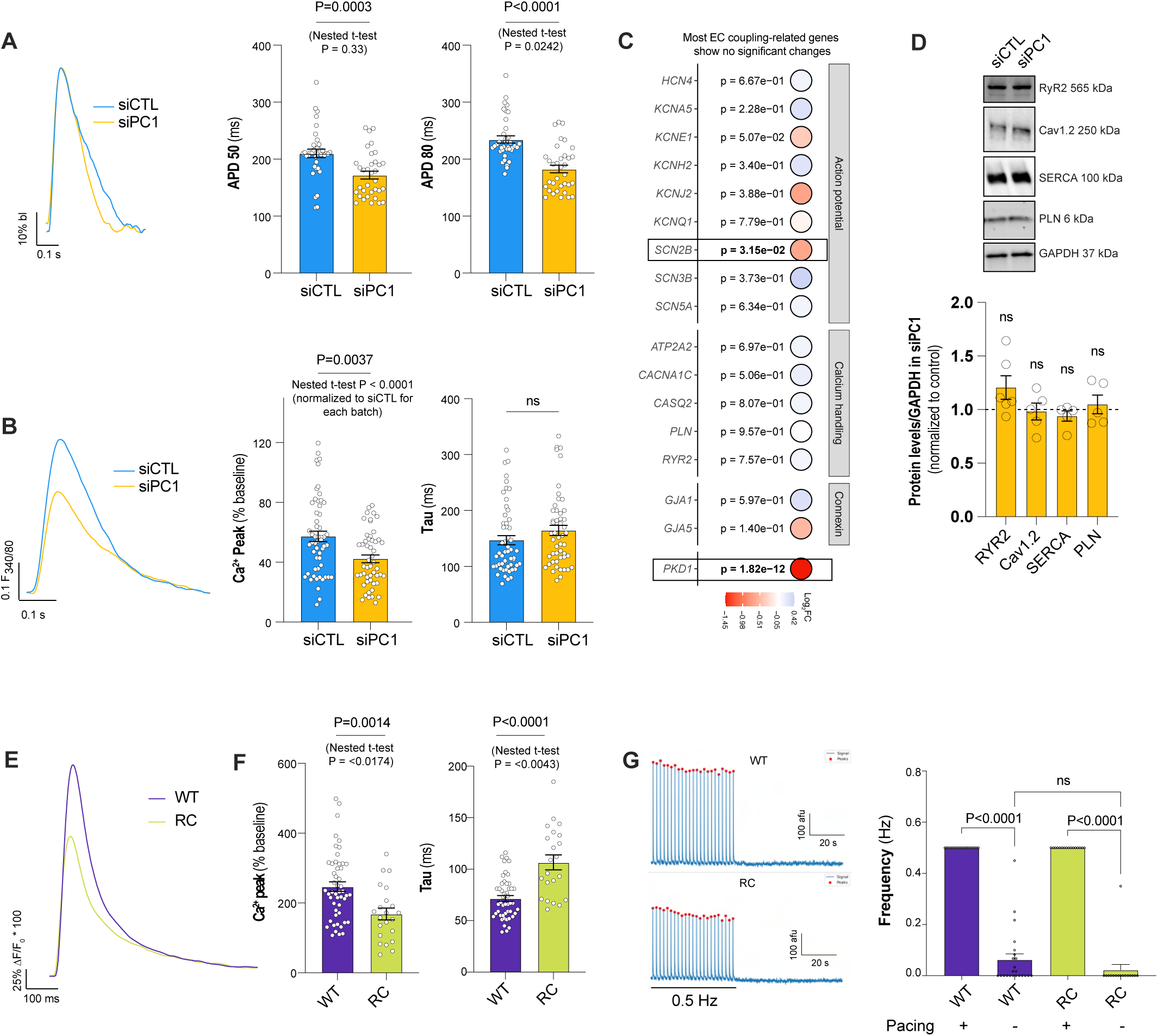
PC1 deficiency regulates excitation-contraction coupling in atrial cardiomyocytes. **A.** Action potential duration was evaluated using FluoVolt and pacing at 2 Hz in hiPSC-aCMs treated with **CTL** and **PC1** siRNA. Shorter action potential duration was observed at 50 and 80% of repolarization in PC1-deficient CMs. **B**. Ca^2+^ cycling was evaluated in hiPSC-aCM and pacing at 2 Hz. A reduced Ca^2+^ transient amplitude was observed after PC1 silencing. **C**. Dot plot shows relevant genes associated with action potential, Ca^2+^ handling, and intercellular communication. No major changes were observed in PC1-deficient iPSC-aCM. **D**. Western blot showing normal levels of RyR2, Cav1.2, SERCA and PLN expression in siCTL and siPC1. **E.** Representative Ca^2+^ transients in **RC** atrial cardiomyocytes, showing reduced Ca^2+^ calcium peak and increased tau compared to **WT** counterparts (**F**). **G**. Representative traces of spontaneous Ca^2+^ events after 1 min of pacing at 0.5 Hz. Quantification shows signal spikes detected using a 10% over baseline threshold during pacing and resting in WT and RC mice. Data are presented as the mean value ± SEM, and the individual data points are also shown. Nested t-tests were used to calculate statistics across multiple batches of cells. Table S1a provides detailed information about the statistical analysis.

Because alterations in APD could lead to impaired Ca^2+^ transients, we assessed Ca^2+^ transients elicited by field electrical stimulation in hiPSC-aCMs. Transient PC1 knockdown reduced Ca^2+^ transients in hiPSC-aCMs (Figure 7B). We investigated the gene expression of key components involved in excitation-contraction coupling, however, we did not observe any significant effect in gene expression or protein levels (Figure 7C-D). These findings agree with our previous report showing that PC1 deficiency shortens APD duration and impinges on Ca^2+^ handling via regulation of Kv channels in ventricular cardiomyocytes^30^, which could explain observed changes in atrial cardiomyocytes (refer to discussion).

We evaluated Ca^2+^ handling in mouse atrial adult cardiomyocytes isolated from WT and RC mice. We found that the peak amplitude of the Ca^2+^ transient induced by electrical stimulation was lower in RC mice compared to WT mice (Figure 7E-F). We then tested whether RC atrial cardiomyocytes displayed spontaneous activity after pacing by measuring Ca^2+^ spikes after stopping the pacing (Figure 7G). We did not observe any differences in the frequency of spontaneous Ca^2+^ events between genotypes. This suggests that the occurrence of arrhythmias in RC mice is not caused by spontaneous activity and may occur at the tissue level under stress conditions (e.g. tachypacing or programmed electrical stimulation).

### Impaired PC1 glycosylation in left atrial appendages from patients with persistent AF

To broaden the translational impact of our findings, we evaluated PC1 levels in left atrial appendage samples obtained from patients in sinus rhythm and persistent AF (without ADPKD, Table S6). Although PC1 total protein levels remain unchanged between samples, we found that the ratio between PC1 top/bottom bands was less abundant in AF compared with SR samples (Figure 8), suggesting impaired PC1 maturation. To confirm the identity of PC1 bands in our atrial samples, we removed immature glycans with EndoH and all glycans with PNGase F. We validated PC1 bands based on molecular weight shift, as has been extensively reported for PC1 studies.^50,65^ These results mirror what has been observed for PC1 maturation in the RC mice, which leads to impaired PC1 trafficking to the plasma membrane.^50^ The differences in PC1 N-terminus molecular size primarily stem from its glycosylation status, which reveals its maturation and trafficking through ER-Golgi-primary cilia in renal epithelial cells. The level of functional PC1 has often been evaluated by assessing its glycosylation status; impaired glycosylation results in immature PC1 that is unable to traffic to the plasma membrane (reviewed in ^65^). These findings suggest that alterations in PC1 may extend beyond ADPKD and likely contribute to AF progression.

**Figure 8:**
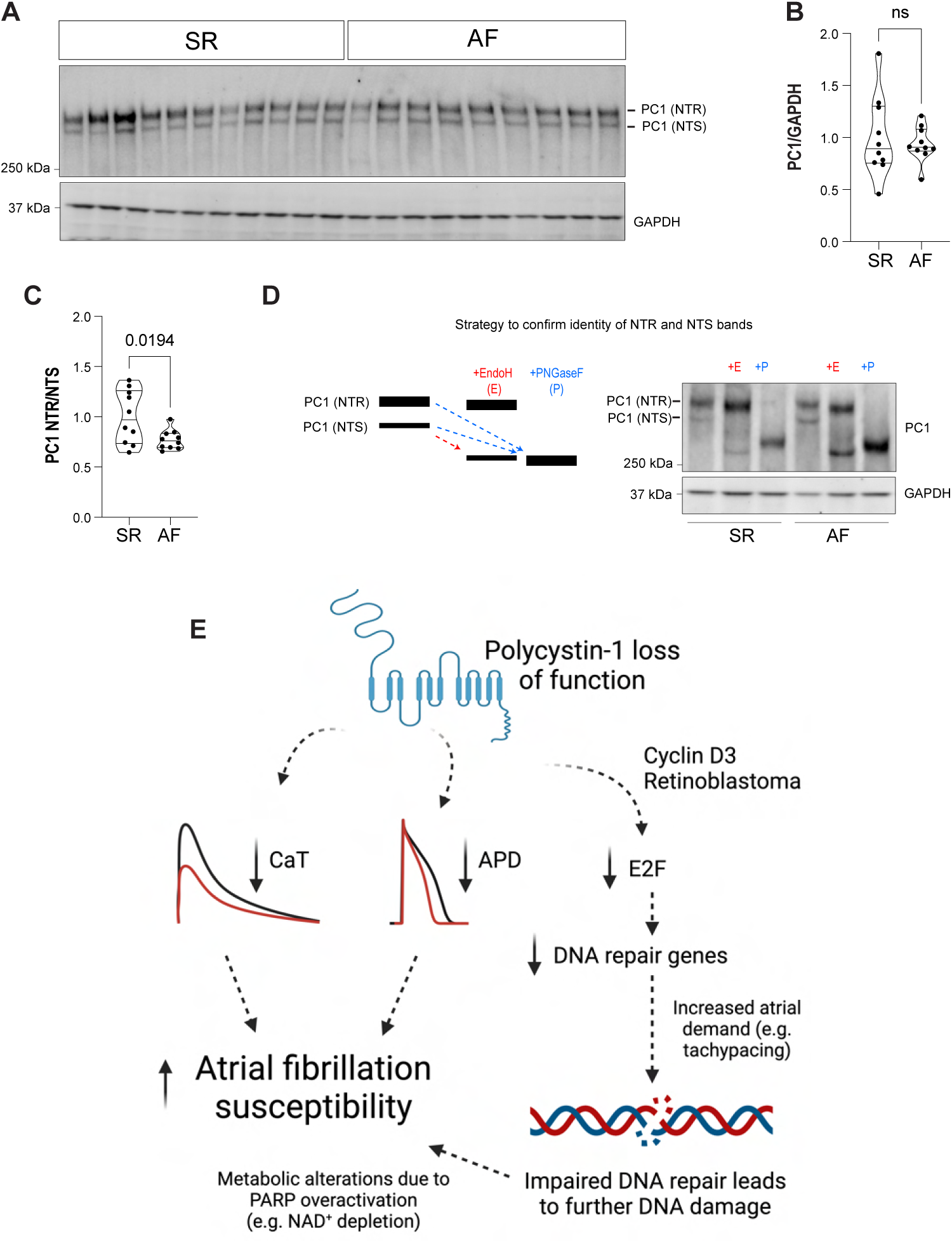
Atrial appendages from patients with persistent AF (AF) exhibit impaired PC1 maturation. Left atrial appendages from 10 control (sinus rhythm, SR) and 10 AF patients were homogenized using protein solubilization buffer (Bio-Rad; Urea, Thiourea, CHAPS buffer), ceramic beads, and a bead homogenizer. **A**. Lysates were separated using Tris-acetate gels (3-8%), and proteins were then transferred to PVDF membranes. **B**. No changes in total levels of PC1/GAPDH were observed among samples (both bands). However, we noted that upper PC1 bands were upregulated in SR vs AF samples. The top band represents the fully glycosylated N-terminal of PC1 (mature; EndoH resistant; NTR), and the bottom band represents the N-terminal of PC1 with immature glycans as found in ER-resident proteins (immature; EndoH sensitive; NTS). **C**. Quantification of PC1 top/bottom bands from blot presented in **A** reveals impaired maturation in AF samples vs. SR. **D**. Representative Endo H and PNGase F showing PC1 immature and mature glycosylation. As expected, mature PC1 (top band) was significantly reduced in AF samples. N = patients are presented as violin plots, including individual data points, median and upper and lower quartiles. No data was excluded or added after the initial analysis design (refer to Table S1a for further statistical information). **E**. Proposed model: loss of function of PC1 reduces DNA repair genes through impaired E2F signaling. This reduction increases susceptibility to DNA damage. Additionally, impaired PC1 leads to shortened action potential and reduced Ca^2+^ transients. These combined factors likely contribute to the increased AF inducibility observed in PC1-deficient mice.

## Discussion

In this study, we demonstrate how PC1 plays a protective role against AF using multiple approaches as follows: 1) PC1 mutations increase *in vivo* AF susceptibility in a mouse model that recapitulates human ADPKD and in cardiomyocyte-specific PC1 KO when compared to their respective controls; 2) Optical mapping in superfused atrial tissue from RC mice revealed circuits for local reentry and impaired response to pacing. *3)* PC1-deficient hiPSC-aCM monolayers exhibit arrhythmic behavior and spiral-wave reentry and phase singularities after tachypacing; *4)* transcriptional profiling identified PC1 as a regulator of key genes involved in DNA repair likely via its C-terminal tail; *5)* loss of PC1 increases susceptibility to DNA damage, which results in arrhythmias; 6) PARP1 inhibition or NR supplementation, strategies aiming to ameliorate NAD^+^ depletion caused by excessive DNA damage, decrease arrhythmias in PC1-deficient monolayers; *7)* Finally, we show that patients with persistent AF have impaired maturation of PC1 protein. A proposed model for increased AF vulnerability in PC1-deficient mice is presented in Figure 8E.

### Role of PC1 in atrial cardiomyocyte DNA repair

Recently, up-regulation of DNA damage responses has been described in the kidneys of ADPKD patients.^66^ DNA damage has also emerged as a driver for AF.^11,37^ AF typically occurs with aging, which may be due to increased levels of DNA damage contributing to further exacerbated underlying EC coupling alterations in AF patients.^6,9–11,67–70^ Increased DNA damage in AF depletes NAD^+^, subsequently impairs mitochondrial function, and increases ROS – a futile cycle that increases AF vulnerability.^11^ A stepwise increase in DNA damage in patients with paroxysmal and persistent AF suggests that DNA damage increases over time and contributes to AF progression.^37^

Our RNA-seq data allowed us to identify that PC1 regulates the expression of DNA repair genes. We subjected our cardiomyocytes to DNA damage stress through tachypacing and etoposide and found impaired DNA repair, leading to further DNA damage in PC1-deficient cells. Interestingly, electrical signals and Ca^2+^ wave propagation were severely impaired after tachypacing and etoposide, which suggests that DNA damage could initiate arrhythmic events in these cells. Indeed, strategies to mitigate metabolic consequences of DNA damage (NR and PARPi) decreased arrhythmia vulnerability in PC1-deficient atrial cardiomyocytes.

Alterations in DNA damage/repair should ultimately lead to changes in excitation-contraction coupling to trigger AF. While this link has been proposed in the literature, the exact connection between DNA damage and irregular electrical activity in the atria remains unclear.^11,37^ DNA damage has been shown to induce Ca^2+^ oscillations in neurons via PARP^71^, and recent studies suggest that DNA damage can activate AMP-activated protein kinase (AMPK) and Ca^2+^/calmodulin-dependent protein kinase (CaMK) to regulate cytosolic Ca^2+^.^72,73^ More studies are needed to dissect how impaired DNA repair increases spontaneous activity in PC1-deficient atrial cardiomyocytes to sustain AF.

### PC1-dependent regulation of atrial EC coupling

APD shortening can cause AF through a shortening of the atrial refractory period.^74–77^ The cardiomyocyte refractory period prevents the firing of ectopic action potentials and helps maintain normal sinus rhythm. Alterations in multiple channels/transporters contribute to the pathogenesis of AF. Interestingly, the gain-of-function of voltage-gated potassium channels (Kv1.5 and Kv4.3) and other channels have been associated with the development of AF because they promote faster repolarization, reducing the refractory period.^78–82^ Our previous studies showed that the human version of PC1 negatively regulates macroscopic currents of human Kv4.3 and Kv1.5^30^, the predominant isoforms leading to atrial repolarization. Consequently, the observed shortening of APD and FPD in hiPSC-aCM following PC1 silencing could likely be attributed to increased Kv4.3 and Kv1.5 currents.

Another major mechanism underlying AF is alterations in Ca^2+^ handling.^74–77^ Our data shows that PC1 deficiency reduced the Ca^2+^ transient peak elicited by pacing. Shortening of APD in PC1-deficient cardiomyocytes could affect Ca^2+^ handling likely through two mechanisms: 1) shorter APD reduces L-type Ca^2+^ channel-Ryanodine receptor 2 (LTCC-RyR2) coupling^83^, and 2) decreases Ca^2+^ entry, which is required to support SR Ca^2+^ refilling.^84^ Further studies are needed to dissect the mechanism driving alterations in excitation-contraction coupling and its potential interplay with DNA damage. Because adult mouse atrial cardiomyocytes isolated from RC mice did not exhibit spontaneous activity after pacing, arrhythmic events may initiate at the tissue level rather than from alterations in individual cardiomyocytes.

### Expanding the scope for PC1 actions beyond ADPKD

Our work challenges the dogma that renal failure increases the occurrence of cardiovascular events in ADPKD patients. When we exposed cKO mice to burst pacing, we found that they had increased rates of AF at similar rates compared with other AF-susceptible mouse models.^39,85,86^ We also provide extensive evidence that PC1 directly regulates atrial cardiomyocyte function. Furthermore, when we looked at a preclinical model for ADPKD at time points that precede renal dysfunction, we also found similar rates of AF susceptibility compared to control mice.

Despite the traditional view that links cardiovascular alterations to renal complications in ADPKD, multiple studies indicate subclinical cardiovascular disease in patients with normal renal function. Indeed, ADPKD patients exhibit subclinical systolic dysfunction, measured as LV strain, before the onset of renal failure.^22,25,26^ Additionally, a higher frequency of congenital heart disease was found, exclusively in patients with *PKD1* pathogenic variants.^23^ Vascular alterations are also present, even in children, before the onset of renal failure.^24^

Increased AF inducibility in young, healthy RC mice before the onset of renal failure is a remarkable finding, suggesting that PC1 alterations increase vulnerability to AF under stress conditions. Demonstrating AF susceptibility before the onset of renal failure underscores the potential impact of cardiac complications in the early stages of the disease, emphasizing the need for early detection and intervention in ADPKD patients (as now is being proposed using LV strain for ventricular function). However, we do not assert that ADPKD patients inherently have a higher propensity for AF before the onset of renal failure, but this possibility has not been studied. Given that a third of total AF cases are asymptomatic^87^, this is a relevant consideration. Literature indicates the prevalence of AF in ADPKD ^21,23,27–29^, and a recent study from the US-National Inpatient Sample database, including 71,531 patients, revealed that a higher risk of AF and atrial flutter was observed among ADPKD in the younger age group (18-39 years of age) compared to those with CKD and without ADPKD (odds ratio of 5.71).^27^

Here, we tested whether PC1 loss of function increases susceptibility to experimental AF induced using programmed electrical stimulation, a well-established method.^88–90^ Our research shows that young cKO mice are susceptible to AF, but this susceptibility may be secondary to ventricular dysfunction, as these mice also exhibit mild systolic and profound diastolic dysfunction, as described.^30^ However, the decline in cardiac function does not progress between 2-4 months of age, and mice do not exhibit ventricular remodeling within this time frame.^31^ On the contrary, in RC mice, we did not observe changes in diastolic function measured by pulse wave and tissue Doppler (E/e’) and only observed a mild reduction in fractional shortening. Based on normal ventricular dimensions obtained by echo, normal histology, absence of fibrosis, and normal connexin expression, there is no evidence of adverse cardiac remodeling in young RC mice. This indicates that the increased susceptibility to AF in RC mice is not a result of ventricular remodeling or increased filling pressure. Our *in vitro* work supports a direct effect of PC1 in atrial cardiomyocytes, as it shows *in vitro* arrhythmias after tachypacing in PC1-deficient hiPSC-aCMs.

Our work also raises the possibility that impaired PC1 signaling could be associated with all-cause AF. In that regard, we found less mature PC1 in human left atrial appendages from patients with persistent AF, and this represents an important area for future investigation. This initial observation requires additional validation as it could be influenced by confounding factors, given the extensive remodeling observed in atrial appendages from AF patients.

Recent findings highlight a higher-than-expected frequency of *PKD1* mutations in seemingly healthy individuals.^15^ This suggests that these mutations may predispose patients to disease rather than causing it, emphasizing the complex nature of ADPKD progression. While it is well-established that hypertension exacerbates ADPKD,^91^ the impact of other cardiac alterations, such as AF, should also be considered. These observations raise the possibility that *PKD1* mutations may have systemic effects, contributing to multi-organ dysfunction and potentially leading to conditions like AF ^21,23,27–29^, even in the absence of kidney disease and vice-versa. Therefore, the role of *PKD1* mutations and their potential influence on AF progression warrant further investigation.

### Study limitations

Our work is relevant because cardiovascular mortality in patients with renal stage disease is 500-1000x higher^92^, and there is a stepwise increase in AF incidence with any reduction in renal function.^14^ This is the first study that provides extensive characterization of the molecular mechanism contributing to atrial fibrillation in a relevant mouse model harboring a relevant mutation for the most common genetic cause of end-stage renal disease. Our findings in young mice are particularly significant as they suggest that the early cardiovascular alterations observed could contribute to the overall progression of ADPKD. However, our work also has limitations because AF susceptibility increased under stress conditions (e.g., programmed electrical stimulation and tachypacing), and spontaneous AF was not assessed in this study. It is highly possible that multiple hits are required to increase vulnerability to AF in ADPKD patients.

Interactions between PC1 and polycystin-2 (PC2, the protein product of *PKD2*) are complex and poorly understood in cardiomyocytes and were not evaluated in the present study. PC2 can exist alone and has independent functions, as a trimer (PC2 3) or tetramer (PC2 4) with itself or as a PC1/2 tetramer at a 1:3 stoichiometry (PC1/2 1:3).^93,94^ There is ongoing debate in the literature regarding the nature of PC2, as it has been shown to function as a non-selective cation channel, potentially operating more like a potassium channel.^95^ Depending on their association, PC1/PC2 (1:3) and PC2 (4) complexes exhibit different functional properties.^94^ Furthermore, the role of PKD1/2 as Ca^2+^ channels in primary cilia remains controversial.^96^ Increased pressure in experimental settings might have disrupted the ciliary structure, leading to Ca^2+^ entry and complicating the interpretation of their function.^97^ Moreover, PC1 fragments have been shown in mitochondria and nuclei and associated with actomyosin complex and other ion channels.^51,54,98,99^ Our data shows no effect of PC2 on Ca^2+^ transients compared to control in hiPSC-aCM (data not shown), suggesting these proteins have different functions in atrial cardiomyocytes. However, more studies are needed to understand the complex nature of polycystins.

Although mouse models have been extensively used to discover AF mechanisms, they are limited by short AF episodes and electrophysiological differences compared to humans. Future studies in large animals could validate findings, but these models also have limitations. Recently, a model for ADPKD with mutant *PKD1* was developed in monkeys^100^ and could be used to determine whether PC1 predisposes to atrial arrhythmias. To address some limitations, we included extensive data from hiPSC-aCM (PC1 knockdown and overexpression) and human samples from patients with AF. The use of hiPSC-aCM in cardiac electrophysiology research is also complicated by the inherent immaturity, however, recent publications have shown hiPSC-aCM recapitulate key aspects of human AF ^101–103^. To mitigate these concerns, we used state-of-the-art differentiation protocols^42,103^ and cross-validated our findings with mouse model samples and the widely used HL-1 cell line.

In recent years, there has been a debate about the effectiveness of animal models in translational research. While animal models have provided valuable insights into fundamental pathophysiology, differences between mice and humans hinder clinical translation. FDA Modernization Act 2.0 has opened the door for alternative approaches to discover new therapies, with the ultimate goal of reducing dependence on animal models that often result in dead ends. An integrated approach, combining animal models with new technologies, such as hiPSC-aCM, is essential for producing rigorous, reproducible studies that can be translated across species. Using this approach, we observed similar phenotypes in all models utilized – human and murine cardiomyocytes, hiPSC-aCMs, and HL-1 cells. Nevertheless, further studies are required to validate these findings and test the impact of multiple human *PKD1* variants on atrial function and AF susceptibility.

### Conclusions

Our work provides evidence that loss of PC1 increases AF inducibility through impaired responses to DNA damage.

## Acknowledgments

We thank *Dr. Peter C. Harris* and the *Mayo PKD Center*, Division of Nephrology and Hypertension, Mayo Clinic, for sharing the RC mouse model. *Dr. Joseph A. Hill,* for the initial discussion of this project and for sharing the adenovirus encoding PC1-CT. Dr. *Robert S. Ross* for his insightful comments to improve this manuscript. Dr. *Akanksha Agrawal* for preliminary analyses. We also thank Dr. Christopher Ward and the Kansas PKD Research and Translational Core Center for sharing the PC1 161F monoclonal antibody.

## Supplemental Material

This section contains:

- Supplemental Methods (references included in the main reference list)
- Supplemental Figure Legends S1-S11
- Supplemental Figures S1-S11
- Supplemental Table Legends S1-S6 (Supplemental Tables attached as spreadsheets)
- Supplemental Video Legends 1-5 (Supplemental Videos attached in mp4 format)

## Supplemental Methods

### Animal Models

All animal experiments were performed in accordance with the National Institutes of Health (NIH) guidelines and approved by the Institutional Animal Care and Use Committee of Houston Methodist Research Institute. Mice were housed with a 12h light/dark cycle with food and water *ad libitum*. The cardiomyocyte-specific PC1 KO (cKO) mouse model (αMhc-Cre;*Pkd1^f/f^*) was generated crossing heterozygous *αMhc*-Cre mice (C57BL/6; Jackson Lab: #011038) with *Pkd1^f/f^* mice (C57BL/6; Jackson Lab: #010671). Pups were born at expected Mendelian ratios and genotyped via PCR analysis of tail snip DNA for the *Pkd1* floxed allele (primers 5’-CCGCTGTGTCTCAGTGCCTG and 5’-CAAGAGGGCTTTTCTTGCTG) and the Cre allele (primers 5’ GATTTCGACCAGGTTCGTTC and 5’GCTAACCAGCGTTTTCGTTC).

Recombination was assessed in isolated cardiomyocytes using the following primers: 5’-CCGCTGTGTCTCAGTGCCTG and 5’-GGTAGCACATAGAAGTGGGTGGC. *Pkd1*^RC/RC^ mouse line was obtained from Dr. Peter Harris at Mayo Clinic, MN, USA. WT controls were generated using a *Pkd1*^RC/+^ x *Pkd1*^RC/+^ breeding strategy, and pups were born at expected Mendelian ratios. The RC (p.R3277C) mutation destroys the *Hinfl* site resulting in a 719-bp PCR fragment spanning the RC allele. Pups were genotyped for this fragment (WT, 439 bp, 280 bp; Pkd1 p.R3277C, 719 bp). Primer used 5’-CAA AGG TCT GGG TGA TAA CTG GTG −3’ and 5’-CAG GAC AGC CAA ATA GAC AGG G-3’. Both male and female mice were used for experiments (2-4 months of age). Refer to Figure S1 for additional information.

### Intracardiac electrophysiology and programmed electrical stimulation

Intracardiac electrophysiology studies were performed on mice as previously described.^38–41^ Animals were anesthetized, and body temperature was maintained using a heating pad. ECG electrodes were inserted into corresponding limbs. The right jugular vein was accessed using blunt dissection, and a longitudinal incision was performed on the jugular vein. A 1.1F octopolar catheter (Millar Instruments) was advanced into the right atrium and ventricle. Proper catheter position is verified by visualization of the waveforms of the intracardiac electrograms of the apex of the right ventricle and the right atrium. The electrode within the right atrium was then used for stimulation. AF susceptibility was then assessed using 3 rounds of 2-second bursts for each cycle with an initial cycle length of 40 ms and decrementing cycle length by 2 ms for a final cycle length of 10 ms. Thirty seconds of recovery were allowed between each burst pacing. AF was defined as disorganized high-frequency atrial signals and the disappearance of P-wave in surface ECG. A smaller number of events were atrial flutter (multiple P-waves with regular atrial signals). Data were analyzed using LabChart (AD Instruments) as previously described.^38–41^ Poincaré plots displaying RR variability were calculated using LabChart during the challenge with the longest arrhythmia. If no arrhythmia was observed, the RR variability was calculated during the last challenge. These plots are meant to demonstrate the variability of RR intervals during induced AF episodes. For P-wave return time, we measured the interval from the end of tachypacing until the first QRS complex, accompanied by clear P-wave, atrial, and ventricular signals. The longest P-wave return time is reported per mouse. For mice without arrhythmias, the time reflects the period for the ECG to return to baseline after programmed electrical stimulation.

### Optical mapping

Mouse atrial tissues were loaded using a modified Langendorff perfusion system. After cannulation, hearts were perfused with oxygenated Tyrode’s solution at 37°C, and tissue health was monitored via ECG. The composition of Tyrode’s solution included 135 mM NaCl, 5 mM KCl, 1.8 mM CaCl_2_, 1 mM MgCl_2_, 10 mM Glucose, and 10 mM Hepes, buffered to pH 7.4. Tissues were loaded for 25 min with 5 µM Rhod-2 AM, 5 µM Blebbistatin, 0.01% pluronic acid in Tyrode’s solution, followed by 10 minutes with 10 µM RH-237. Subsequently, the left atrial appendage was dissected, spread on a polydimethylsiloxane (PDMS) surface, and secured with insect pins. Point electrical stimulation was delivered via platinum wire electrodes. Atrial tissues received biphasic pulses of 3 V for 5 ms each pulse.

Optical mapping was performed using a MiCaM Ultima system (100×100 pixel camera) with LEX2 530 nm LED illumination equipped with a 5x objective lens (51.9 μm/pixel). During recordings, the peristaltic flow of Tyrode’s solution was stopped to reduce motion artifacts. Atrial tissues were paced at 6-12 Hz for 30 seconds to achieve steady-state conditions before recording 1024 frames at 500 Hz. Data analysis was executed using MATLAB, with frequency filtering above 50 Hz, Gaussian spatial filtering (radius=5), and temporal median filtering (5 points). Baseline drifts were corrected using a fourth-degree polynomial fit. Data was normalized so the values for each pixel range from 0 to 1. TIF and AVI files were exported using batch processing for further analysis and arrhythmia evaluation. Activation maps were generated using Electromap, focusing on the depolarization midpoint of action potentials. Additionally, Fast Fourier Transform (FFT) analysis was used to examine dominant frequency. A moving average filter (window size=9) was used to reduce noise and calculation errors using a Python script and the scipy.fft package. Python scripts were optimized with the assistance of OpenAI GPT-4 and validated by comparing output results to the ones obtained with Rhythm or ElectroMap. For experiments in iPSC-aCM, cells were loaded with 10 µM Rhod-2 AM at 37°C for 15 minutes, followed by a 30-minute de-esterification at room temperature. A 0.63X objective lens was used to collect fluorescence from monolayers (133.3 μm/pixel). Point stimulation was delivered using a custom-made 3D printed insert made of PLA with a pair of platinum wires. Pulses were sequentially delivered at 2-6 Hz (5V, 5 ms, biphasic) for 1 minute to reach steady-state levels before recording 1024 frames at 200 Hz. Signals were processed as described above. Phase mapping was calculated using the Hilbert transform, centered to adjust for data skewness, using a Python script and the scipy.signal.hilbert package.

Experiments in HL-1 monolayers were recorded using a MiCAM Ultima imaging system (SciMedia) attached to a trinocular camera port in an Olympus IX73 microscope with a 10X objective. Cells were loaded with Fluo-8AM (10 μM) in Tyrode’s buffer for 25 min followed by an additional 25 min for de-esterification and imaged in the presence of 100 μM Norepinephrine (the concentration in Claycomb’s media). HL-1 monolayers were imaged at room temperature. Spontaneous activity was recorded at 200 fps for 20 seconds. Data were exported and analyzed using Rhythm 1.2 software and Matlab as previously described.^104^

### Cellular Models

Human episomal iPSC cell line (A18945, hPSCreg Name TMOi001-A) was purchased from Thermo Fisher. Cells were cultured and maintained on Matrigel (Corning) in mTeSR plus (STEMCELL Technologies) or E8 (Thermo Fisher) media according to the manufacturer’s protocol. Differentiation to hiPSC-aCMs was performed at 90% confluence using a previously published protocol.^105^ Cells were differentiated in cardiomyocyte differentiation media: CDM3 or RPMI 1640 media supplemented with B27 Minus Insulin. CDM3 is a mixture of RPMI 1640 media supplemented with 500 μg/ml O. sativa-derived recombinant human albumin and 213 μg/ml l-ascorbic acid 2-phosphate. Differentiation started with CHIR99021 (6μM, Selleckchem) in cardiomyocyte differentiation media for 2 days. Then, the media was replaced and supplemented with 2 μm Wnt-C59 (Tocris). On day 3, retinoic acid (Tocris) was spiked for a final concentration of 1μm. On day 4, the media was changed and supplemented with 1μm retinoic acid. After day 6, media was switched to RPMI supplemented with B27 until cells started beating (days 9-10). Selection media (RPMI media without glucose supplemented with B27) was used to metabolically select atrial cardiomyocytes. Cells were then passaged onto new Matrigel (Corning) coated plates with RPMI + B27 media until experiments were performed after day 35 of differentiation. Refer to Figure S9 for further information.

HL-1 cells were obtained from Sigma Aldrich (SCC065) and cultured in Claycomb media (10% FBS, 10 μM Norepinephrine, 2 mM L-Glutamine, 1X Penicillin/Streptomycin in Claycomb Basal Medium) on 0.02% gelatin (EMD Millipore) and 5 μg/ml fibronectin (Sigma) with daily media changes until experimentation as previously described.^43,44^

### siRNA knockdown

hiPSC-aCMs were transfected with control (SIC001, Universal Control, Mission Sigma), PC1 (SASI_Hs01_00136684/PKD1, sequence #1, SASI_Hs01_0013-6686/PKD1, sequence #2, Mission Sigma) or PC2 (SASI_Hs01_0010-0551/PKD2, Mission Sigma) siRNAs using Lipofectamine RNAiMAX (Thermo Fisher) into wells containing RPMI B27 media (75 nM siRNA final concentration). HL-1 cells were transfected using control (SIC001, Universal Control, Mission Sigma) or PC1 (SASI_Mm01_0014_9980, Mission siRNA) siRNA into wells containing Claycomb media as described above.

### Multielectrode Arrays

MEA plates with carbon nanotube electrodes (8×8 grid, 150µm electrode x-y separation, Matrigel-coated 6-well plates, Alpha Med Scientific) were sterilized using 70% ethanol and UV light, followed by washes with sterile water, and coated with Matrigel (1:100 dilution). hiPSC-aCMs were plated onto carbon nanotube electrodes at a density of ∼150,000 cells/cm^2^. Cells were allowed to recover and synchronize for ∼7 days and then transfected with control and PC1 siRNA. Electrical activity was recorded after 48 h at a sampling rate of 3 KHz using a MED64 Presto (Alpha Med Scientific) at 37°C. Monolayers exhibited perfect synchronization under spontaneous conditions. Frequency and FPD were calculated using Symphony software (Alpha Med Scientific). Time stamps for each electrode’s beats were exported and analyzed using an R code (R Studio 2023.03.0 Build 386/R version 4.2.3). In order to conduct an unbiased analysis, an R script was employed to process all monolayer data. The script was optimized with the assistance of OpenAI GPT-4 and validated by using synthetic data. Furthermore, the data were manually inspected to ensure the accuracy of the measurements. The main R additional packages for analysis were tidyverse (includes dplyr), fields, and reshape2. Other packages, such as ggplot2, RColorBrewer, ggpubr, were primarily used for data visualization, while gridExtra, Cairo, and pdftools were used for working with images and graphics. The average frequency for each electrode and the entire monolayer was used to separate beats. A normal beating pattern was considered when all active electrodes had electrical signals within 10 ms (most beats propagate within this period). Ectopic activity (second electrical signal within interbeat interval) and propagation-delayed beats (maximum time of propagation >10 ms) were calculated as well. In addition, we calculated the number of electrodes with intermittent activity (skipped beats). In addition, conduction velocity was calculated using linear regression, and surface plots of a thin plate spline fit the data. An arrhythmic monolayer was defined by ectopic activity, delayed propagation, and/or monolayers with conduction block between adjacent electrodes.

### Adult mouse atrial cardiomyocyte isolation

WT and RC mice (2-4 months old) were anesthetized using 5% isoflurane in 100% O_2_ prior to euthanasia by cervical dislocation. The heart was quickly dissected, and the aorta was cannulated. Isolated hearts were perfused with Solution 1 (130 mM NaCl, 5 mM KCl, KH2PO4 0.5 mM, and 10 mM glucose, EDTA 5 mM, 30 mM taurine, 10 mM 2,3-butanedione monoxime, 10 mM HEPES, pH 7.4) for 2 minutes. Then, the perfusion line was switched to Solution 2 (same composition as Solution 1 without EDTA and supplemented with 1 mM MgCl_2_, 0.025 mg/mL Liberase TM, and 0.025% Trypsin). Atria were excised, dissociated by pipetting, and filtered using a 250 µm filter. Ca^2+^ was then introduced stepwise up to 1.8 mM, then plated on Matrigel-coated 35 mm glass bottom dishes. Only quiescent atrial cardiomyocytes with elongated structures and defined striations were used in this study.

### Measurement of action potentials

hiPSC-aCMs were loaded with FluoVolt for 25 min at room temperature in Tyrode’s buffer (135 mM NaCl, 5 mM KCl, 1.8 mM CaCl_2_, 1 mM MgCl_2_, 10 mM HEPES, 5.6 mM Glucose, pH 7.4) per manufacturer’s protocol. Cells were washed and imaged in Tyrode’s at 37°C using an IonOptix imaging system equipped with a 470 LED, capturing at an acquisition rate of 250 frames per second. Electrical field stimulation (5 ms 10V biphasic pulse) at a frequency of 2 Hz was delivered using Myopacer (IonOptix) and a custom dish insert, with platinum wires approximately 8 mm apart. Data were analyzed using IonWizard software (IonOptix), with an average taken from at least 5 action potentials during steady-state conditions.

### Intracellular Ca^2+^ measurements

hiPSC-aCMs were loaded with Fura-2AM (5 μM) for 25 min, washed, and further incubated for an additional 25 min to promote de-esterification in Tyrode’s buffer, all at room temperature. Cells were then imaged and paced using an IonOptix imaging system as described above. Fluorescence detection was conducted at 250 Hz, alternating between 340 nm and 380 nm LED illumination, synchronized with IonOptix. We upgraded our IonOptix imaging system with 470 LED and emission cubes to detect Fluo-4 fluorescence during our studies. Hence, isolated WT and RC adult mouse atrial cardiomyocytes and HL-1 cells were loaded with Fluo-4AM (5 μM) using the same loading protocol described above for Fura-2AM. Ca^2+^ transients in adult cells were evaluated at a frequency of 1 Hz, whereas spontaneous activity HL1 was evaluated in HL1 (as described in the limitation section of this manuscript Discussion section). Data were analyzed using IonWizard software (IonOptix), with an average taken from at least 5 Ca^2+^ transients during steady-state conditions.

### Assessment of SR Ca^2+^ stores

hiPSC-aCMs were loaded with Fluo-4AM as described above and paced at 2 Hz, and SR Ca^2+^ stores were assessed based on previously described methods.^30^ The electrical stimulation was stopped, and caffeine (10 mM) was quickly perfused for 10 seconds. The amplitude of the caffeine-induced Ca^2+^ transient was used to estimate SR Ca^2+^ stores.

### Western Blots

Proteins were resolved in 4-20% TGX under reducing and denaturing conditions. Proteins were then transferred to nitrocellulose membranes using a Trans-blot turbo blotting system (Bio-Rad). Membranes were then blocked with 5% milk in Tris-buffered saline with 0.05% Tween (TBST). Monoclonal 7e12 antibody against PC1 N-terminus (Santa Cruz, sc-130554, 1:100) was used for 2 hours at room temperature, while all other antibodies were incubated overnight at 4°C at the following dilutions Chk1 (Santa Cruz sc8408, 1:100), Rad51 (Cell Signaling, 8875s, 1:1000), TOP2a (Cell Signaling #12286, 1:1000), CEP55 (Cell Signaling, 81693s, 1:1000), E2F2 (Invitrogen, PA5-77950, 1:1000), PC2 (Santa Cruz, sc-47734, 1:100), Phospho-Rb (Ser780) (Cell Signaling, #8180, 1:1000), Rb (4H1) (Cell Signaling, #9309, 1:1000), hFAB™ Rhodamine Anti-GAPDH #12004167 (1:5000).

Secondary antibodies Dylight 800 (Anti-rabbit or anti-mouse, Cell Signaling), and StarBright 700 (Anti-rabbit or anti-mouse, Bio-Rad) were used for detection using ChemiDoc MP (Bio-Rad). In some experiments, to increase the signal-to-noise ratio, a biotinylated secondary antibody (anti-mouse or anti-rabbit depending on the primary antibody, Cell Signaling), streptavidin HRP (Cell Signaling) and ECL ClarityMax (Bio-Rad) were used. Chemidoc detected saturated pixels during exposure, and non-saturated images were analyzed using ImageLab (Bio-Rad).

### Processing of human left atrial appendages

Tissue (∼50 mg) was homogenized using chaotropic total protein solubilization buffer that contains NDSB 201, urea, thiourea, and CHAPS (Bio-Rad) supplemented with 1x protease and phosphatase inhibitors (Cell Signaling). The tissues were cut into small pieces and homogenized using a bead mill homogenizer with 2.8mm ceramic beads at room temperature. Samples were spun at 10,000 *xg,* and clear lysates were used for protein concentration determination using RC DC Protein Assay (Bio-Rad). Samples were then stored at −80 °C. For western blot analyses, samples were mixed with XT sample buffer 2X and reducing agent (Bio-Rad) and loaded directly into 3-8% Tris-acetate gels to determine PC1 protein levels as described above. To validate the identity of PC1 bands recognized by the 7e12 monoclonal antibody, we evaluated PC1’s glycosylation status as described.^50,65^ Samples were concentrated using 10 MWCO spin columns to remove chaotropic lysis buffer and resuspended in Cell Signaling Lysis Buffer (containing 1% Triton) with protease and phosphatase inhibitors. EndoH and PNGase F (New England BioLabs) were used to remove immature and mature glycosylation as per manufacturer instructions.

### Adenoviral Overexpression

Adenovirus expressing LacZ or PC1-CT under CMV promoter was previously described^24^. Adenoviruses were amplified in Adeno-X 293 cells (Clontech) until late cytopathic effects. Media and cells were then collected, and 3 cycles of freeze-thaw cycles were performed. Titer was calculated using Adeno-X™ qPCR Titration Kit (Takara Bio). Cells were then transduced using a multiplicity of infection of 10 viral particles per cell, and transduction efficiency was 100%, measured by ZsGreen1 fluorescence (co-expressed under a second CMV promoter; Fig. 6B). Cells were lysed after 48-72 h for RNA studies. Adenovirus expressing mCherry-PC1-CT-V5 was cloned using Adeno-X Adenoviral System 3 (CMV; #632269, Takara).

### Bulk-RNA Sequencing

Total RNA was extracted using Quick-RNA Miniprep (Zymo Research), and concentration was measured at 260/280 nm. RNA samples (∼500 μg, Abs_260/280_ > 2) were sent to GeneWiz for further analysis. To ensure a fully unbiased RNA-seq, we also requested their bioinformatics analysis service. The company’s pipeline includes quality controls of RNA integrity, library preparation, next-generation sequencing, and data analysis. Quality control and RNA integrity were evaluated using Agilent TapeStation 4200 and Qubit by Genewiz before proceeding with RNA-seq. All samples showed an RNA integrity index > 9.5. Further details about RNA processing were provided by Genewiz as follows: *RNA sequencing libraries were prepared using the NEBNext Ultra II RNA Library Prep Kit for Illumina following the manufacturer’s instructions (NEB, Ipswich, MA, USA). Briefly, mRNAs were first enriched with Oligo(dT) beads. Enriched mRNAs were fragmented for 15 minutes at 94 °C. First-strand and second-strand cDNAs were subsequently synthesized. cDNA fragments were end-repaired and adenylated at 3’ends, and universal adapters were ligated to cDNA fragments, followed by index addition and library enrichment by limited-cycle PCR. The sequencing libraries were validated on the Agilent TapeStation (Agilent Technologies), and quantified by using Qubit 2.0 Fluorometer (Invitrogen) as well as by quantitative PCR (KAPA Biosystems)*.

*The sequencing libraries were clustered on 1 flowcell lane. After clustering, the flowcell was loaded on the Illumina HiSeq instrument (4000 or equivalent) according to the manufacturer’s instructions. The samples were sequenced using a 2×150bp Paired-End (PE) configuration. Image analysis and base calling were conducted by the HiSeq Control Software (HCS). Raw sequence data (.bcl files) generated from Illumina HiSeq was converted into fastq files and de-multiplexed using Illumina’s bcl2fastq 2.17 software. One mismatch was allowed for index sequence identification*.

*Data Analysis: after investigating the raw data quality, sequence reads were trimmed to remove possible adapter sequences and nucleotides with poor quality using Trimmomatic v.0.36. The trimmed reads were mapped to the Homo sapiens reference genome available on ENSEMBL using the STAR aligner v.2.5.2b. The STAR aligner is a splice aligner that detects splice junctions and incorporates them to help align the entire read sequences. BAM files were generated as a result of this step. Unique gene hit counts were calculated using feature Counts from the Subread package v.1.5.2. Only unique reads that fall within exon regions were counted*.

*After the extraction of gene hit counts, the gene hit counts table was used for downstream differential expression analysis. Using DESeq2, a comparison of gene expression between the groups of samples was performed. The Wald test was used to generate p-values and log2 fold changes. Multiple comparison correction was achieved using Benjamini Hochberg (p-adjusted)*.

We used iDEP9.5 (http://bioinformatics.sdstate.edu/idep95/) and pyDESeq2 (https://pydeseq2.readthedocs.io/en/latest/index.html) to verify the accuracy of the calculated results and found them to be in alignment GeneWiz’s rlog transformed counts and GO terms. Our lab then further conducted a further analysis of the differentially expressed genes to identify regulated pathways using RStudio and Clusterprofiler 4.0. We followed validated pipelines for over-representation analysis (ORA) and Gene Set Enrichment Analysis (GSEA) (https://yulab-smu.top/biomedical-knowledge-mining-book/index.html).^45^ We used Gene Ontology Biological Processes, KEGG, and MSigDB v7.5.1 databases to investigate the potential actions of PC1 in atrial cardiomyocytes. Additionally, we utilized ingenuity pathway analysis software (IPA, Qiagen) to further explore differentially expressed genes. For ORA and IPA analysis, given that PC1 manipulation had a mild but significant effect on gene expression, we decided to use a cut-off of FDR < 0.05 and an absolute value of log_2_FC > 0.5 to identify affected pathways. Summary bubble plots were generated in R using the ggplot2 package, and circular plots were generated using the GOplot package. Refer to Table S1 for further details.

### Real-time PCR

RNA was isolated using a Quick-RNA microprep kit (Zymo Research). RNA concentration was measured with a Qubit, and 500 ng of RNA was converted to cDNA using iScript Reverse Transcription Supermix (Bio-Rad) as per the manufacturer’s instructions. RNA levels were assessed by semiquantitative PCR using iTaq Universal SYBR Green Supermix (Bio-Rad) with a CFX Connect (Bio-Rad). Relative transcript levels were analyzed using CFX maestro with the comparative Ct method and normalized to four housekeeping genes (GAPDH, HPRT1, TBP, and PGK1). Relative transcript levels were normalized to their own control levels (per batch of cells), averaged, and analyzed using GraphPad Prism software using a two-tailed one-sample t-test. Primers used in this study are as follows:

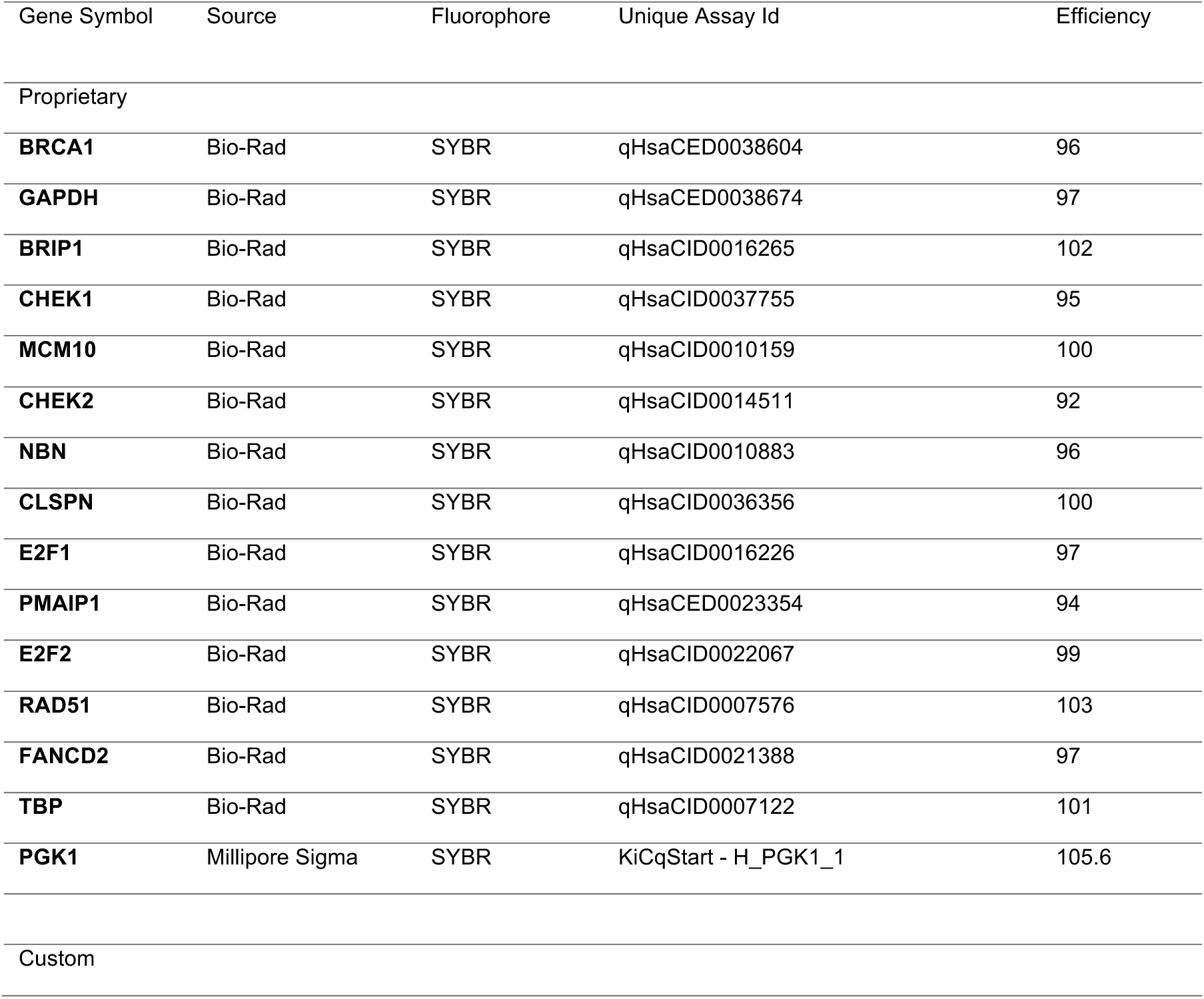
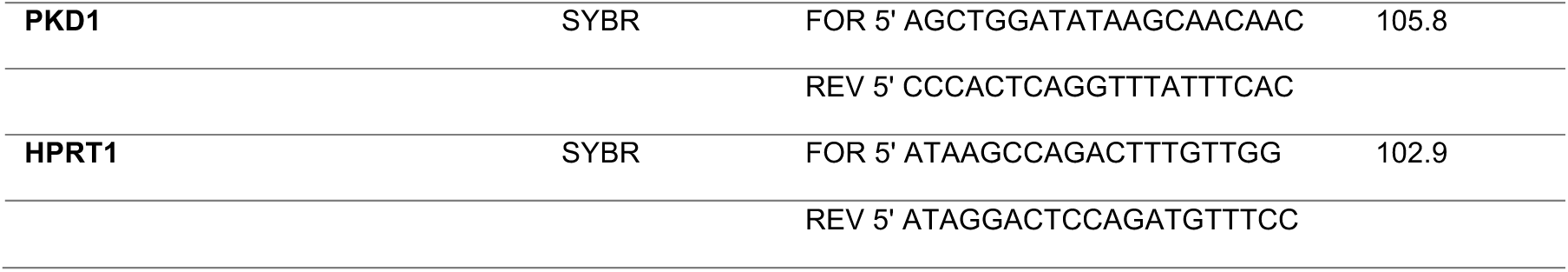

### Immunofluorescence

Cells were washed, fixed with 4% paraformaldehyde, permeabilized with 0.1% Triton X-100, and blocked in 5% normal goat serum in phosphate buffered saline (PBS). Samples were then incubated overnight at 4° C with the following primary antibodies: Cardiac Troponin T (abcam #ab8295, 1:500), α-actinin (Invitrogen #701914,1:200), MLC2a, (Invitrogen #PA6-53041, 1:500), MLC2v (Proteintech, #10906-1-AP, 1:500), Kv1.5 (Sigma Aldrich #P9357 1:500), p-Histone H2A.X (S139) (20E3) (Cell signaling #9718S 1:400), Poly/Mono-ADP Ribose (Cell signaling #83732s 1:1000), mCherry (Cell signaling #5536T, 1:100), V5 (Bio-Rad # MCA1360, 1:100), PC1-CT (recognizes human PC1-CT, Kerafast EJH002, developed by PKD Core, U. Maryland, 1:100), PC1-CT 161F (recognizes human PC1-CT, gift from Dr. Ward, Kansas PKD Core^53^, 1:100). Highly Cross-absorbed anti-IgG (H+L) secondary antibodies conjugated to Alexa Fluor 488, 555 or 633 and DAPI or Hoechst were used for detection. Immunofluorescence were imaged on 40x oil-immersion using an Olympus Fluoview 3000 confocal microscope using diode lasers with excitations at 405 nm, 488 nm, 561 nm, and/or 633 nm using proper emission filters or Cytation 1 with DAPI, FITC and TRITC filters. Bleed-through was monitored using antibody controls and analyzing potential artifactual channel overlap.

In some experiments, hiPSC-aCM monolayers were fixed in cold methanol (−20 C) to measure total cell count and Poly-ADP-Ribose (PAR) levels after optical mapping. Cells were permeabilized with 0.5% Triton X-100 for 15 min, blocked in 5% normal goat serum, and then stained for α-actinin, PAR and DAPI. We captured 15×15 images covering a total surface of 77 cm^2^ (14 mm coverslips = 154 cm2 = 50% monolayer coverage) using a 10X objective and LED cubes to detect DAPI (nuclei), GFP (α-actinin) and TRITC (PAR) wavelengths using a high-throughput microscope (Cytation 1, Agilent). Images were analyzed using Gen5 software (Agilent). A nuclear mask was created using DAPI signals to measure PAR intensity per nuclei. Total cell count and PAR-positive cells were calculated per monolayer.

### Histology

Whole heart and kidneys were dissected from WT and RC mice (2-3 months old), rinsed with PBS, and fixed in 4% paraformaldehyde diluted in PBS for 24 hours. Samples were processed, sectioned, and stained, blinded by the Houston Methodist Research Pathology Core. Paraffin-embedded tissues were analyzed using H&E or Masson’s Trichrome stains. Pictures were taken using an EVOS-FL digital microscope at 4X and stitched together. Images were adjusted using Adobe Photoshop for brightness, contrast, and color balance.

### DNA comet assays

Cells were lifted, resuspended in PBS, and then embedded in low melting agarose on a comet slide at 37 **°**C at a 1/10 v/v ratio (R&D - Single Cell DNA Comet assay kit). Comet assays were performed as per the manufacturer’s protocols. Samples were then immersed in lysis buffer for 60 minutes at 4 **°**C and switched to unwinding buffer (NaOH 200mM, 1 mM EDTA). Alkaline running buffer (same as an unwinding buffer) was used for electrophoresis at 4 **°**C at 20 V for 30 minutes. Samples were washed, fixed with 70% Ethanol, air-dried, stained with SYBR Safe (Thermo Fisher), and imaged using ZOE fluorescent imager (Bio-Rad). Cometscore (http://rexhoover.com/index.php?id=cometscore) was used to measure olive tail moments.

### Tachypacing protocol

Tachypacing was performed using C-Pace (IonOptix) and 6-well carbon electrode plates in hiPSC-aCM. A custom 3D printed 6-well plate-like adapter was used to fit six 35 mm dishes with glass bottom coverslips. The pacing parameters were set at 10V and 2 ms pulses at 8 Hz. To better understand the protective role of PC1 in atrial cardiomyocytes, we opted to stress the monolayer with a higher frequency. It should be noted that hiPSC-aCM can only capture frequencies up to 6 Hz. However, during AF episodes, local high-frequency activity can reach 8 Hz or higher.^106^ Thus, we chose to tachypace at a frequency of 8 Hz as a suitable stress strategy to reveal PC1 protective actions in atrial cardiomyocytes. The temperature was measured in pilot experiments using a thermocouple connected to a Fluke thermometer and remained at ∼37 °C during the entire experiment. Viability was assessed as described in the Results section. To block PARP1 activity, ABT-888 (Veliparib, Tocris) was added to the culture media to a final concentration of 20 μM. Cells were preincubated for 15 min before tachypacing and kept during the entire experiment.

## Supplemental Figures Legends

**Figure S1:**
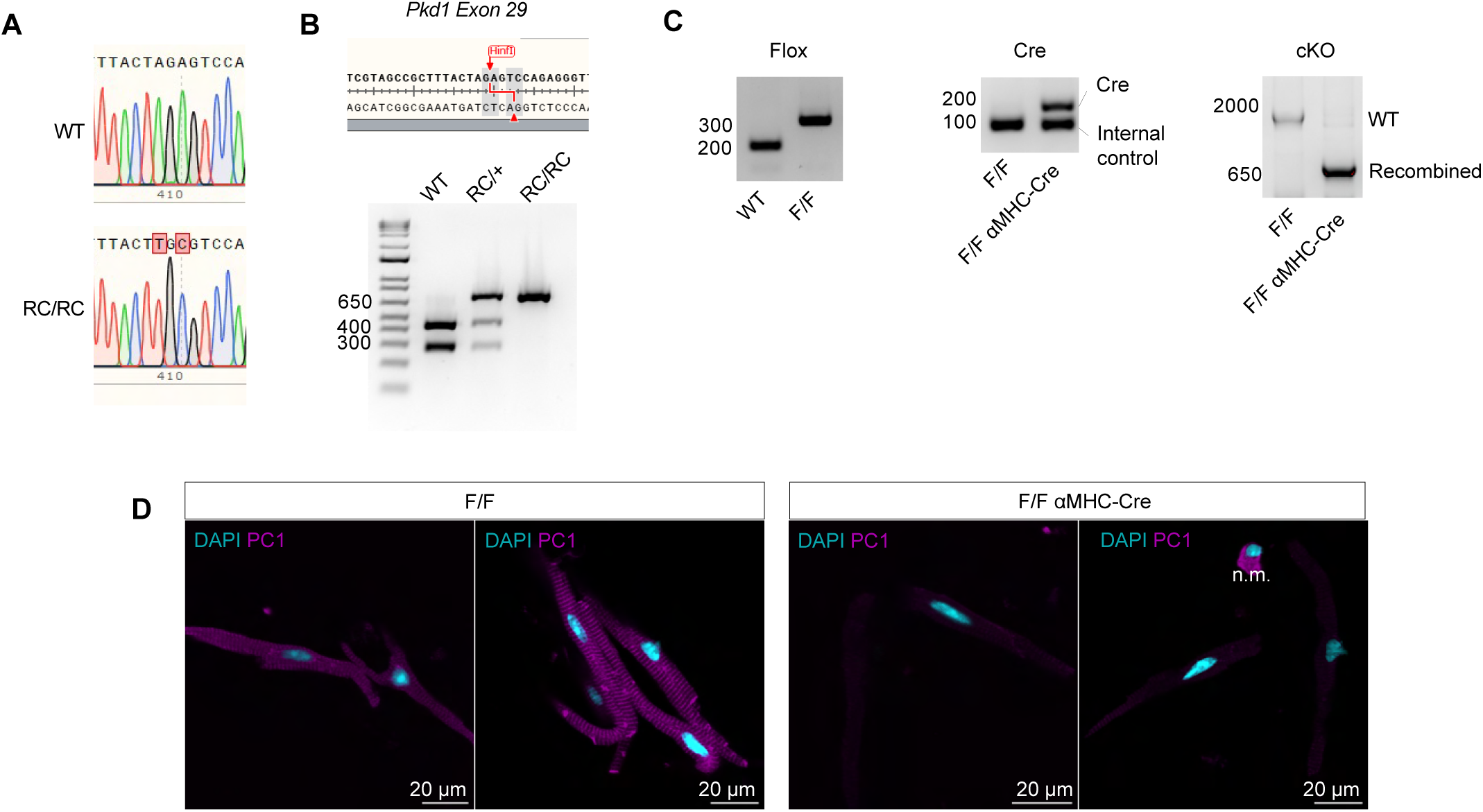
Validation of RC and cKO mouse models. **A.** Sanger sequencing confirmed the point mutation in our RC model. **B.** A PCR reaction was used to amplify the region containing the *Hinfl* restriction enzyme site, which is affected by the R3277C mutation. WT PCR samples were cut by *Hinfl,* resulting in 2 bands, whereas a single band was used to confirm genotyping in RC mice (see Methods). **C.** For cardiomyocyte-specific PC1 KO mice, we detected flox sites in the *Pkd1* gene and αMHC-Cre using PCR in tail samples. *Right panel*, isolated atrial cardiomyocytes were used to demonstrate the recombination of *flox* alleles in the *Pkd1* gene. **D**. Immunostaining in adult mouse atrial cardiomyocytes showing effective deletion of PC1 protein in cardiomyocyte-specific PC1 KO mice.

**Figure S2:**
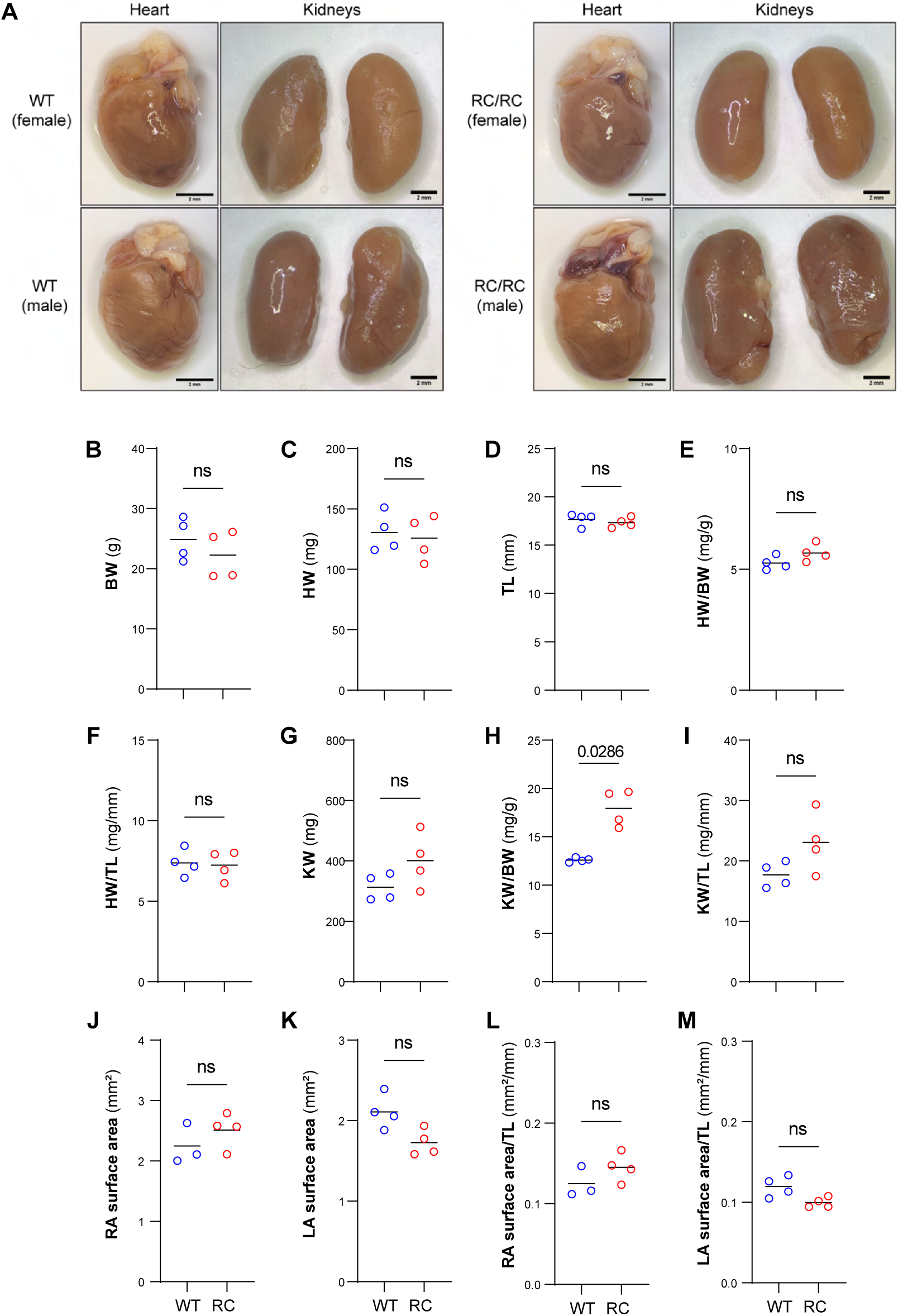
Heart and kidney morphometry in WT and RC mice. **A)** No major difference was observed between WT and RC hearts isolated from mice at 2-3 months of age. Scale bar = 2mm. **B)** Body weight. **C)** Heart Weight. **D)** Tibia length. **E)** Heart weight divided by body weight. **F)** Heart weight divided by tibia length. **G)** Kidney weight. **H)** Kidney weight divided by body. **I)** Kidney weight divided by tibia length. Although a slight but significant increase in KW/BW was observed in RC mice as described in the renal field ^50^, normalization to TL showed more variable results. **J)** Right and **K)** left atria surface area in WT and RC hearts were measured using ImageJ and normalized to tibia length, respectively (**L-M**). Details of statistical tests are presented in Table S1b.

**Figure S3:**
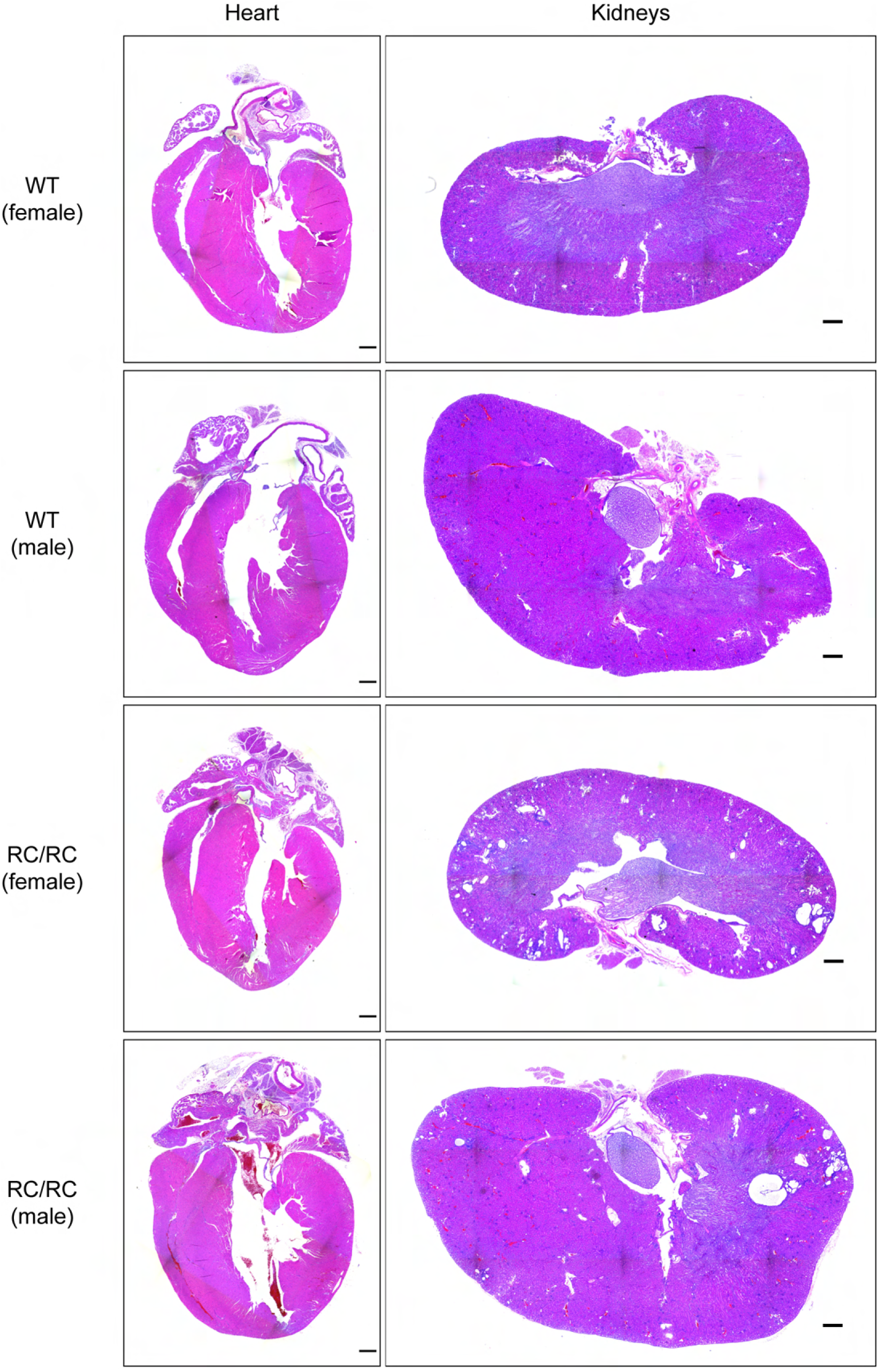
Hematoxylin and eosin staining in heart and kidney sections obtained from WT and RC mice: Longitudinal 4-chamber views of the heart and longitudinal kidney sections were stained using standard hematoxylin and eosin (H&E) techniques. No major alterations in histology were observed in the hearts of males and females. In striking contrast, RC kidneys showed evidence of cystic formation as expected at this age (2-3 months) because cystogenesis develops in utero and progresses to renal failure after 9 months.^50^ Images were taken using a 4X objective and stitched using an EVOS FL imaging system. Scale bar = 500 μm

**Figure S4:**
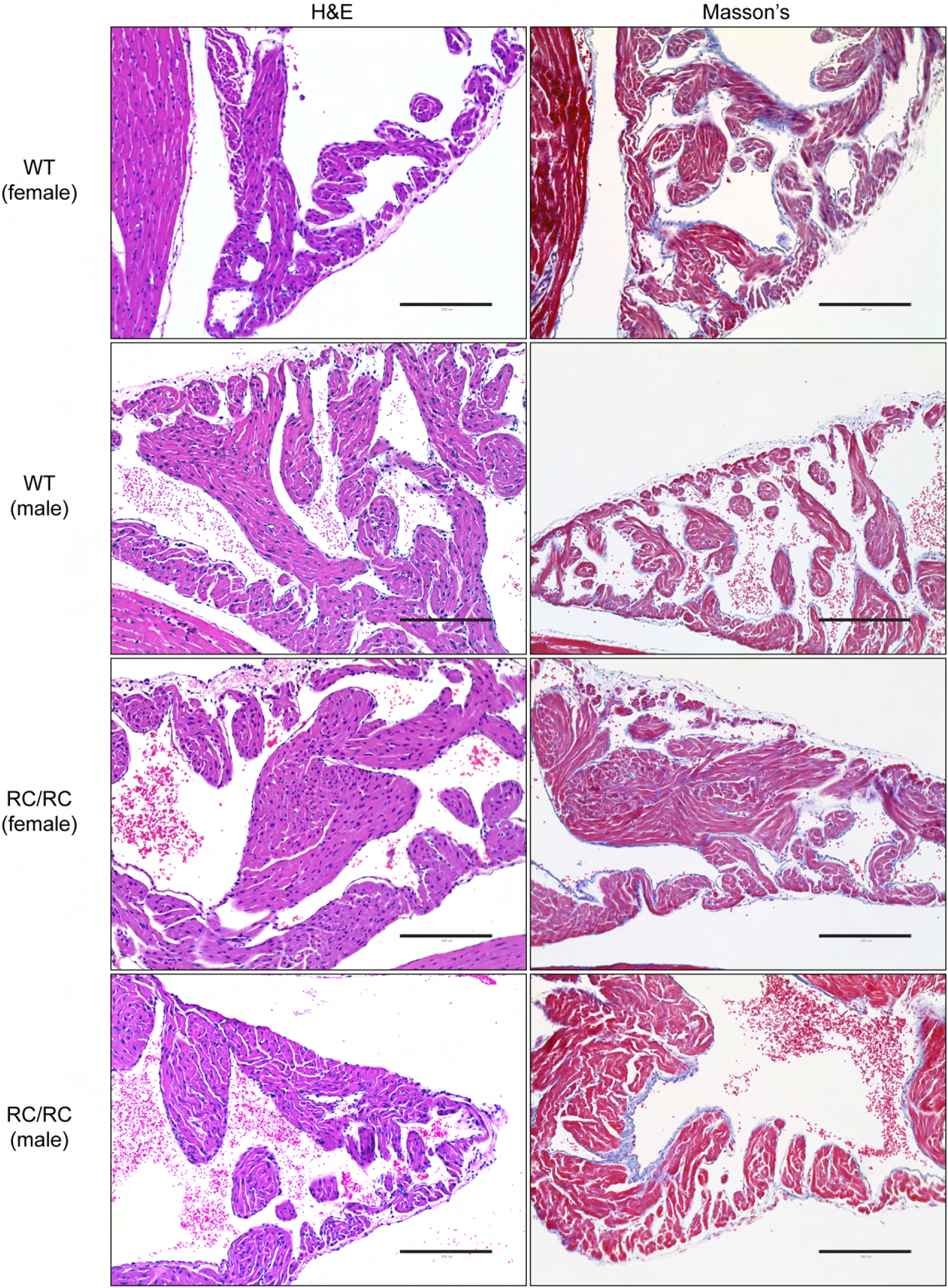
Normal left atria histology in RC mice. 4-chamber views of the heart were stained using H&E and Masson’s trichrome staining. Normal histology was observed. Scale bar = 200 μm

**Figure S5:**
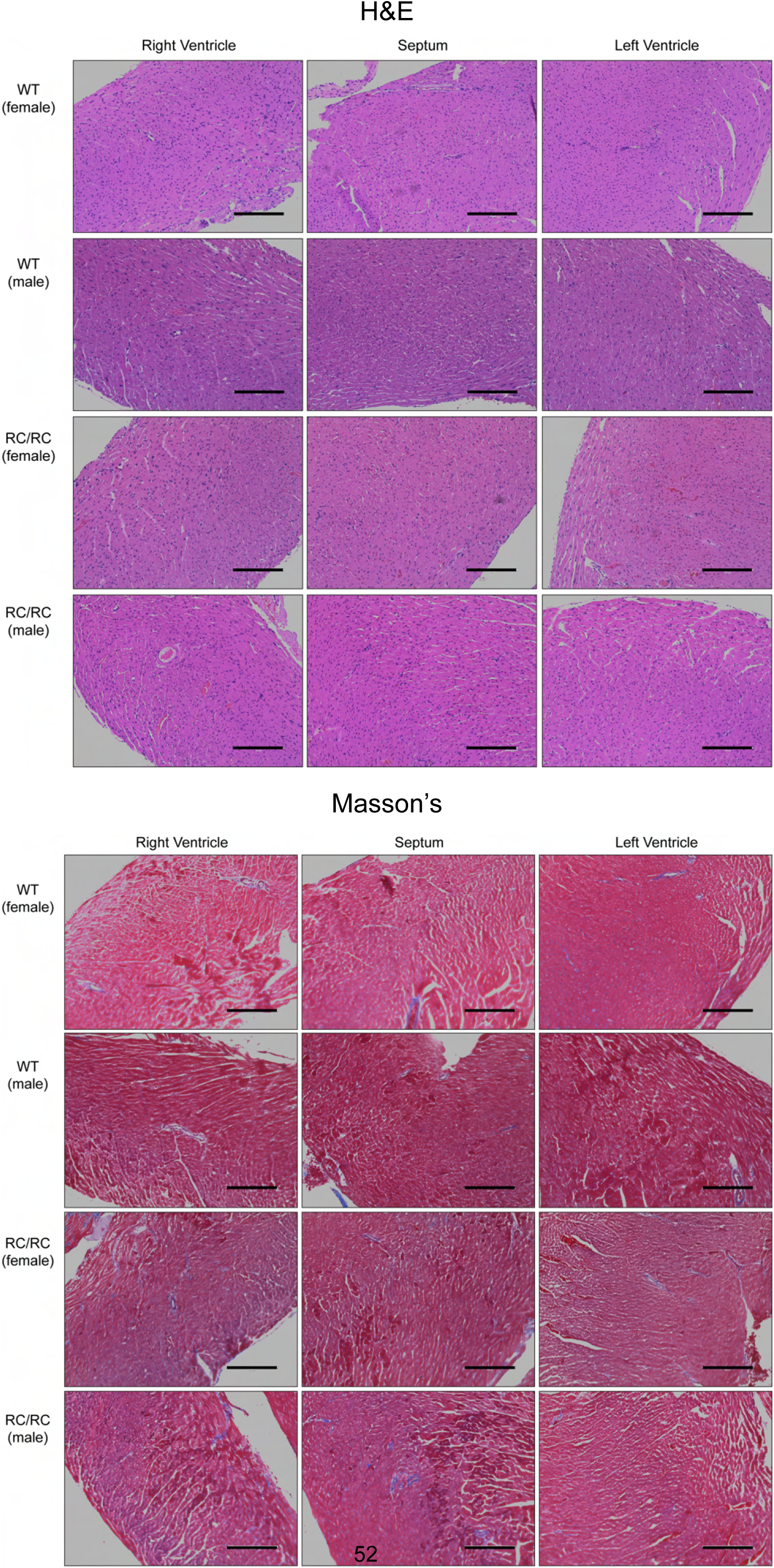
H&E and Masson’s trichrome staining in 4-chamber view ventricles obtained from WT and RC mice. The right ventricle, septum, and left ventricle mid-sections in 4-chamber view were stained. Scale bar = 200 μm

**Figure S6:**
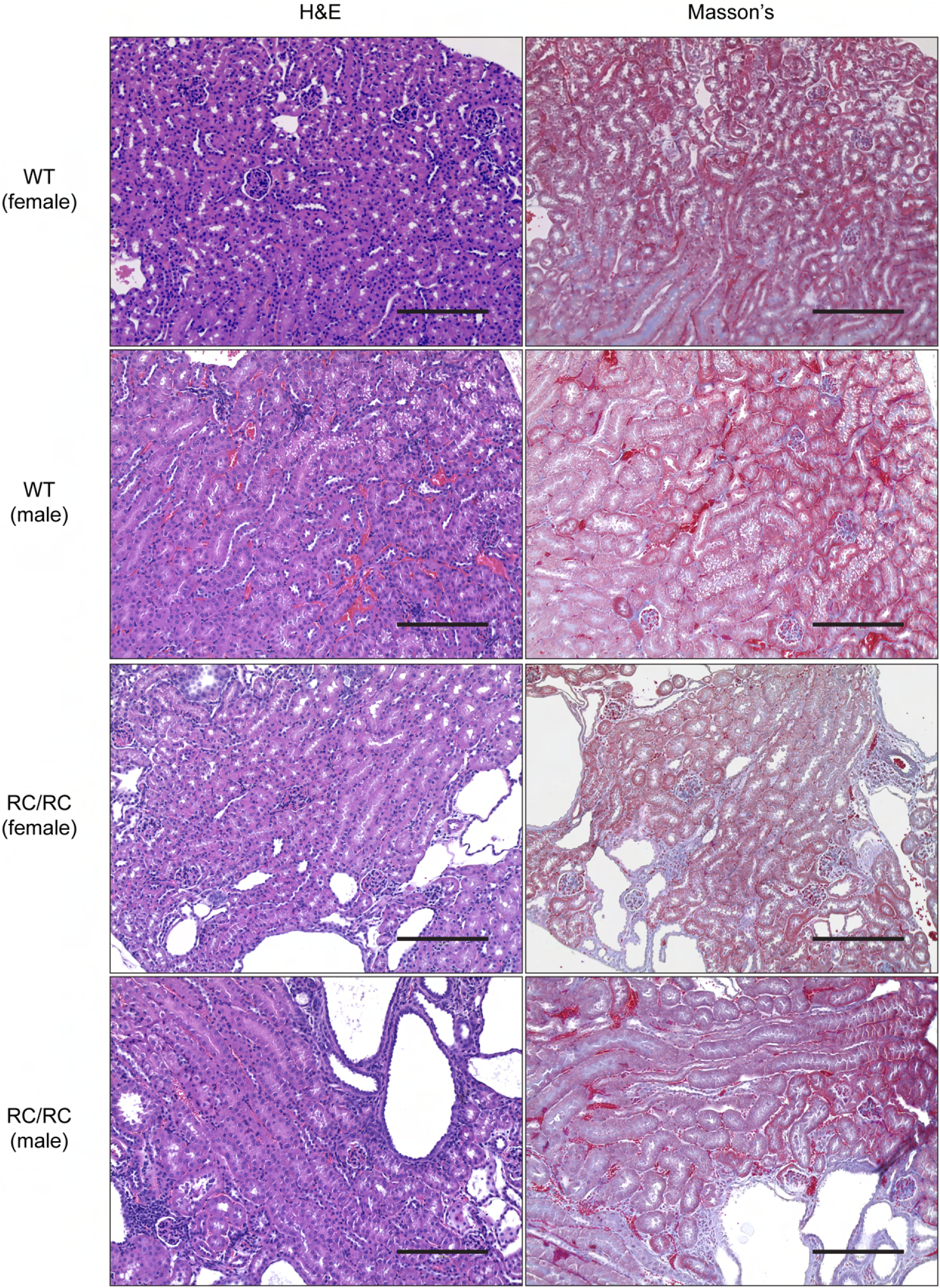
Increased renal fibrosis and cyst formation in RC mice. Longitudinal kidney sections were stained using H&E and Masson’s trichrome staining. Scale bar = 200μm.

**Figure S7:**
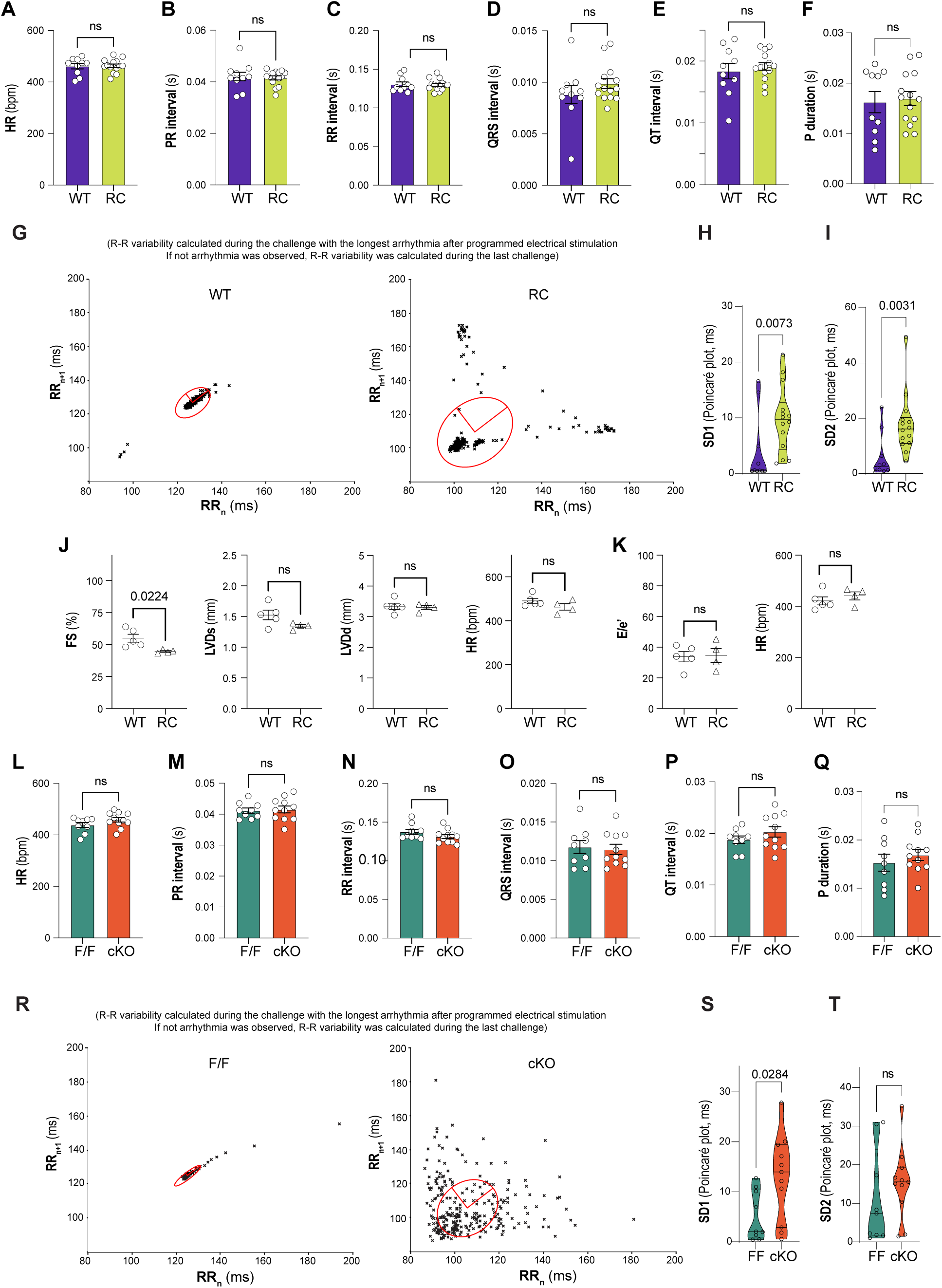
Electrocardiogram parameters and echocardiography reveal normal heart rate and intervals. Mice were anesthetized, and the recording catheter was inserted through the jugular vein. Intracardiac and surface electrocardiograms were used to determine the optimal catheter positioning. Then, animals were maintained until the heart rate stabilized around 425 bpm. The Figure shows baseline ECG data monitored prior to burst pacing and compared in F/F vs. cKO and WT vs. RC. No significant changes in **A.** Heart rate, **B.** PR interval, **C.** RR interval, **D.** QRS interval, and **E.** QT interval were observed. **F**. P wave duration. **G.** Representative Poincaré plots showing RR variability at the longest arrhythmia. If no arrhythmia was observed, we calculated the RR interval in the last challenge. **H** and **I** show Poincaré SD1 and SD2 calculations. Data are presented as violin plots, including individual data points, median and upper and lower quartiles. **J**. Echocardiography shows minimal changes in fractional shortening (FS) and no significant changes in left ventricular dimensions in systole and diastole (LVIDs and LVIDd). HR during measurement is shown in the right panel. **K**. Pulse wave and tissue doppler (E/e’) revealed no changes in diastolic function in RC mice (2-3 months old). HR during measurement is shown in the right panel. ECG data was monitored prior to burst pacing and compared in F/F vs. cKO. No significant changes in **L.** Heart rate, **M.** PR interval, **N.** RR interval, **O.** QRS interval, and **P.** QT interval and **Q**. P wave duration were observed. **R**, **S,** and **T** provide Poincare plots as described in **G-I**. Additional information regarding the statistical analysis can be found in Table S1b.

**Figure S8:**
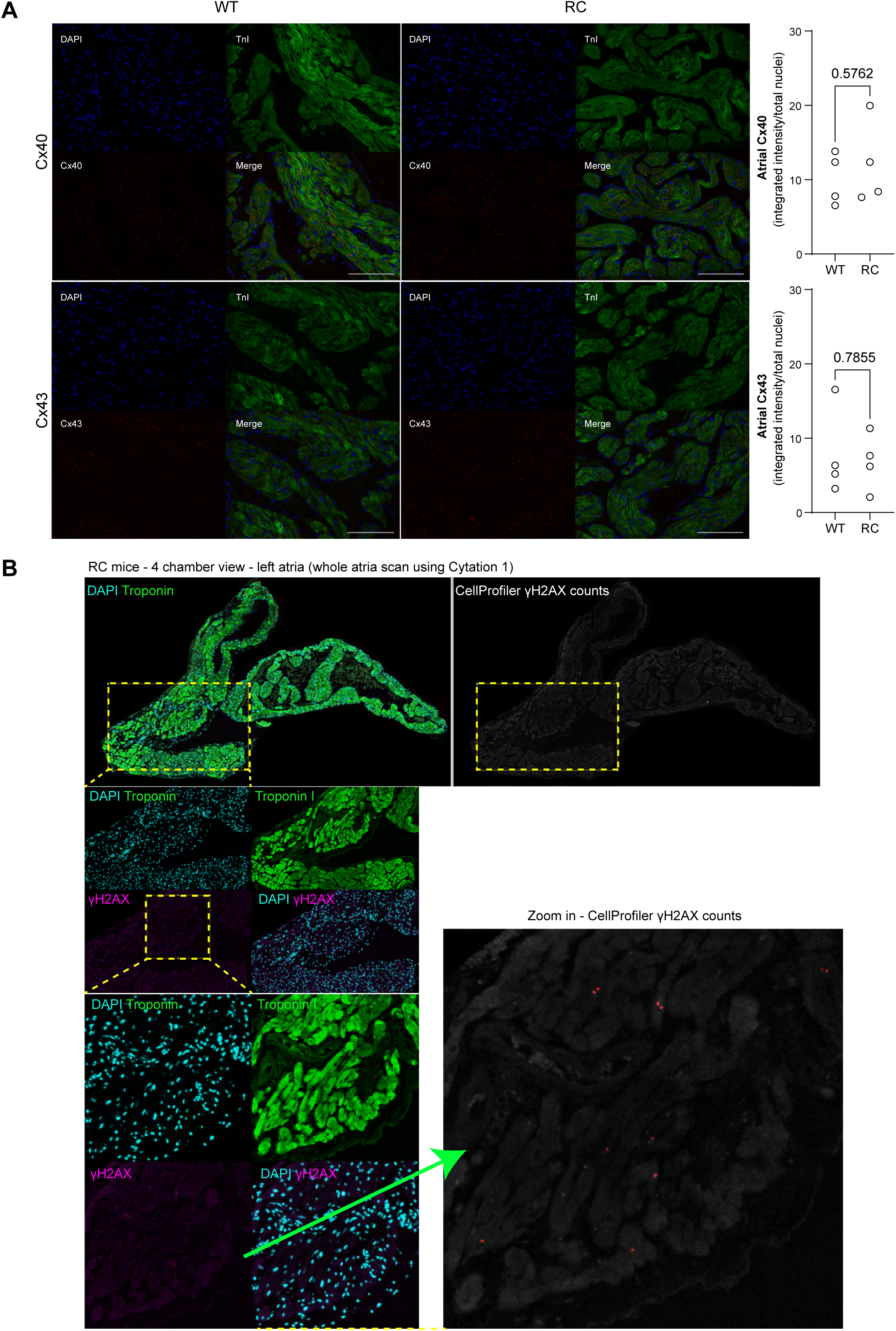
Connexin 40 and 43 immunostaining. **A.** Tissue were stainined for troponin-I and either Cx40 or Cx43. Fluorescence intensity was measured using Cellprofiler and normalized to the number of nuclei. Scale bar = 100 μm. Refer to Table S1b for additional details about the statistical analysis. **B**. Clustering of γH2AX-positive cells in RC left atria. The data presented in this figure was obtained by scanning the entire left atria (4-chamber view) using a 20X objective with Cytation 1. Quantification in Figure 5B was carried out on confocal images randomly captured within the left atria and quantified using CellProfiler.

**Figure S9:**
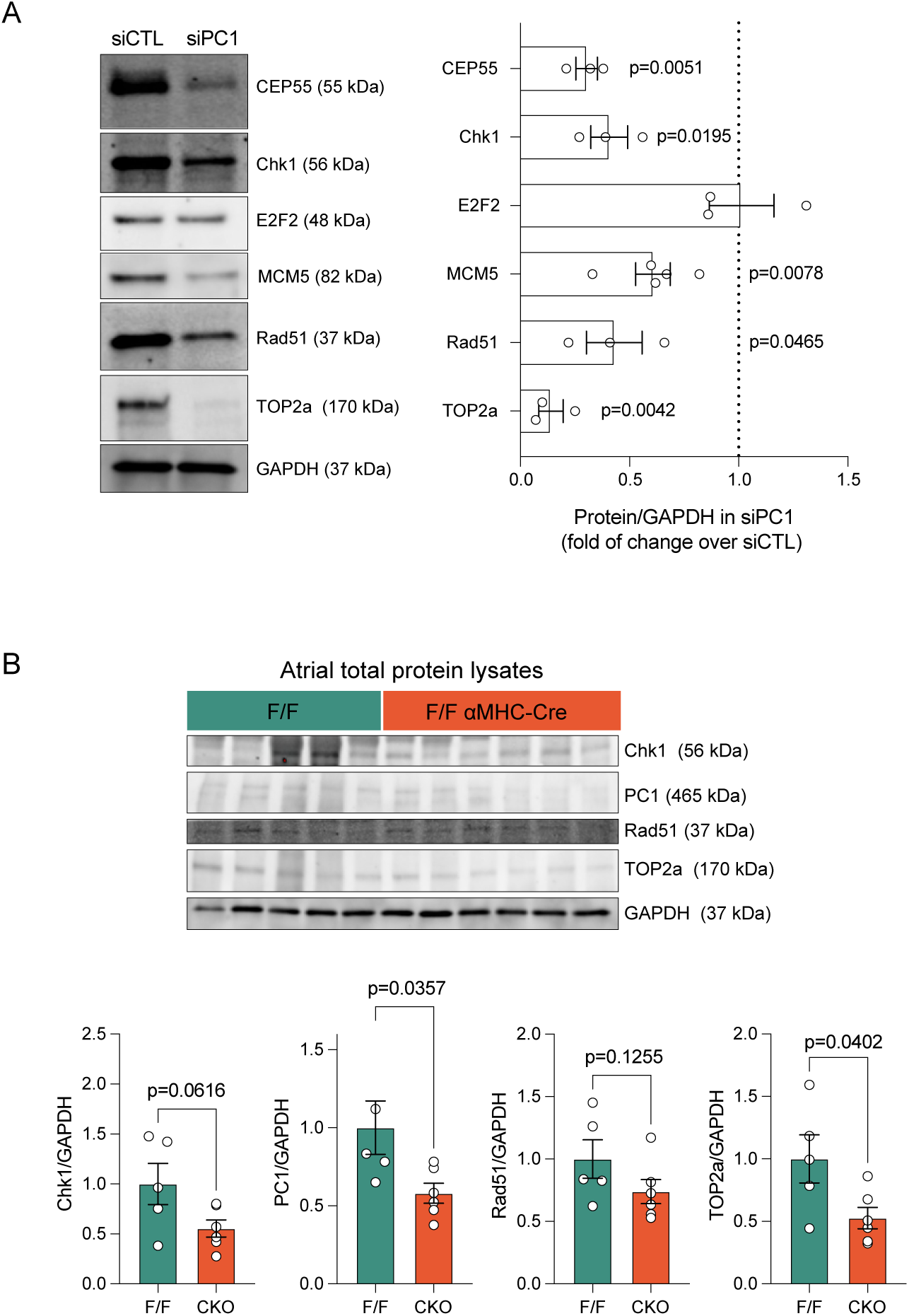
Protein levels following PC1 knockdown in hiPSC-aCMs. **A**. Protein expression levels of interest candidates identified by RNA sequencing were measured by western blot after 48h of PC1 knockdown and compared to control siRNA-treated hiPSC-aCMs. Key regulators involved in DNA repair, such as CEP55, Chk1, E2F2, MCM5, Rad51 and TOP2a were significantly decreased. **B.** Protein levels in F/F and cKO atrial tissue. After the catheterization procedure, the atria were dissected, and total protein lysates were prepared and analyzed using Western blot. GAPDH was used for normalization. Chk1, PC1, Rad51 and TOP2a protein levels normalized by total protein in F/F and cKO mice. Additional information regarding the statistical analysis can be found in Table S1b.

**Figure S10:**
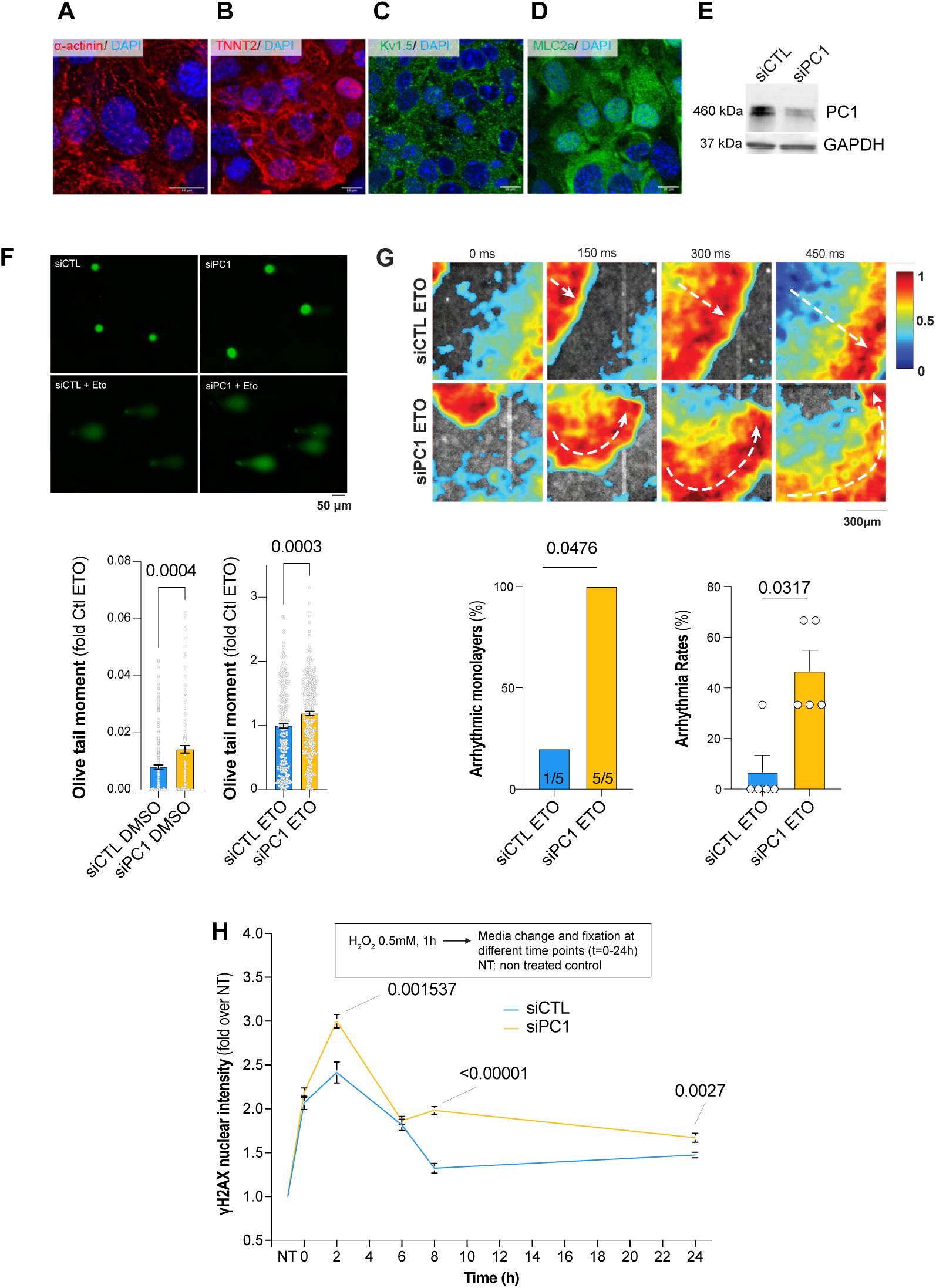
Increased DNA damage and arrhythmias in PC1-deficient HL-1 cells. Immunostaining with cardiomyocyte-specific markers **A.** α-actinin, **B.** Troponin T, **C.** Kv1.5 and **D.** MLC2a. Scale bar = 10 μm. **E.** Efficient PC1 silencing in HL-1 cells at 48h compared to control siRNA. **F.** DNA comets assay reveals increased baseline DNA damage in PC1-deficient cells exacerbated with etoposide treatment (double- and single-stranded breaks are detected when using comet assays in alkaline conditions). **G.** Ca^2+^ optical mapping in HL-1 cells treated with etoposide revealed increased non-linear propagation and spiral-like waves in PC1-deficient cells. **H.** γ-H2AX clearance assays, cells were exposed to hydrogen peroxide for one h, then washed, and then allowed to recover in complete culture media. Non-treated (NT) cells were used as the baseline. Cells were either fixed immediately after damage or within 2-24 h of recovery to track DNA repair (decreases in γ-H2AX nuclear intensity over time as previously described^107^). After fixing, cells were permeabilized and then stained for γ-H2AX. The resulting nuclear levels of γ-H2AX fluorescence intensity are shown. Additional information regarding the statistical analysis can be found in Table S1b.

**Figure S11:**
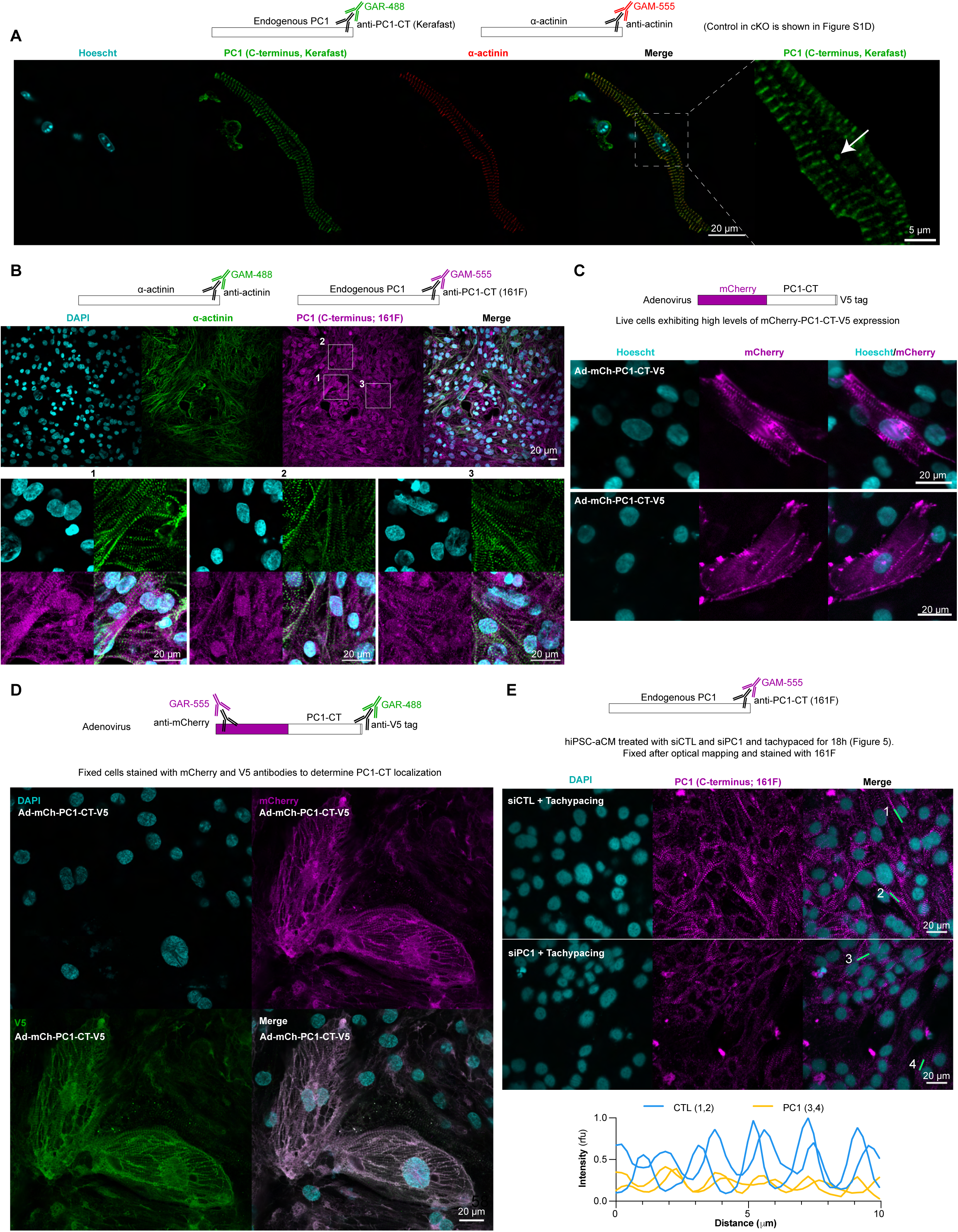
Localization of PC1 in atrial cardiomyocytes. **A**. The Kerafast antibody targeting mouse PC1-CT in adult mouse atrial cardiomyocytes reveals a striated pattern with nuclear puncta (white arrow). **B**. Similar results were observed in hiPSC-aCM stained with the 161F antibody targeting human PC1-CT. **C**. Adenovirus expressing mCherry fused to PC1-CT shows a striated localization with discrete nuclear puncta in live hiPSC-aCM. **D**. Following, fixation, the cells were stained with anti-mCherry and -V5 antibodies, showing striated and nuclear fluorescence. **E**. PC1 staining after tachypacing in both siCTL and siPC1 hiPSC-aCM illustrates an effective knockdown. Line fluorescence intensity profiles for 2 cells are displayed for each condition.

## Supplemental Table legends

**Table S1:** Comprehensive details on the statistical analysis of all Figures.

**Table S2:** Differentially expressed genes in hiPSC-aCM treated with PC1 and CTL siRNA (5 differentiation batches containing a PC1 knockdown and control sample; each one sequenced in duplicate). All genes with an adjusted p-value (FDR) < 0.05 are included in this table. Comparison between groups was performed using DESeq2 (Wald test p-value and Benjamini-Hochberg adjusted p-value). Batch columns show normalized RNA counts. Log2FC: log2 fold of change; Padj: P adjusted; CTL: control; PC1: Polycystin-1 siRNA (refer to Table S1 and Methods for further details).

**Table S3:** Gene ontology Biological Process in PC1-deficient (siPC1) hiPSC-aCM compared to control (siCTL). All genes with an adjusted p-value (FDR) < 0.05 and absolute value of log2 FC > 0.5 were used for these analyses using clusterProfiler v4.0 and IPA. Table S3a: Gene Ontology Biological Process, S3b: IPA analysis (refer to Table S1 and Methods for further details).

**Table S4:** Differentially expressed genes in hiPSC-aCM treated with adenovirus expressing PC1- CT and LacZ (3 biological replicates from the same differentiation batch; each one sequenced in duplicate). All genes with an adjusted p-adj (FDR) < 0.05 are included in this table. Comparison between groups was performed using DESeq2 (Wald test p-value and Benjamini-Hochberg adjusted p-value). Replicate columns show normalized RNA counts. Log2FC: log2 fold of change; Padj: P adjusted; LacZ: adenovirus expressing LacZ; PC1-CT: adenovirus expressing PC1 C-terminus. (refer to Table S1 and Methods for further details).

**Table S5: A.** GSEA analysis in RNA-seq databases using Hallmark pathways and custom gene sets for PC1 expression analysis as described in the main text. (refer to Table S1 and Methods for further details). **B.** GSEA analysis in RNA-seq databases using C2 canonical pathways (refer to Table S1 and Methods for further details).

**Table S6:** Characteristics of patients used for biochemical experiments. Data are mean ± SD. CAD, coronary artery disease; MVD/AVD, mitral/aortic valve disease; LVEF, left ventricular ejection fraction; ACE, angiotensin-converting enzyme; AT, angiotensin receptor.

## Supplemental Video legends

**Supplemental Video 1:** Ca^2+^ wave propagation in the superfused left atrial appendage obtained from WT mice. The tissue was paced at 6-12 Hz using a pair of platinum wires located at the bottom of the tissue. The denoised, filtered, and normalized signal (ranging from 0-1 per pixel) is shown with a blue to red-jet color scale. Real-time quantifications from two regions of interest (ROI 1 and 2; highlighted in top panels) are displayed in top and bottom plots.

**Supplemental Video 2:** Ca^2+^ wave propagation in the superfused left atrial appendage obtained from RC mice as described in Supplemental Video 1.

**Supplemental Video 3:** Optical mapping of Ca^2+^ signals in hiPSC-aCM monolayers after tachypacing. The video revealed spontaneous spiral-wave reentry in siPC1 during point stimulation, whereas siCTL shows unidirectional propagation. *Top panel*, Ca^2+^ wave propagation (denoised, filtered and normalized signal per pixel is shown in blue to red jet scale). *Bottom panel, the* Hilbert transform shows phase singularities in siPC1 calculated with scipy.signal package (jet color scale from -π to +π).

**Supplemental Video 4:** Spline surface plots were generated to visualize spontaneous electrical signal propagation in the 8×8 MEA grid. The video shows 249 consecutive beats obtained from a siCTL hiPSC-aCM monolayer treated with etoposide for 2 h. The time scale is shown on the right side, ranging from 0 to 5 ms, using the viridis color scale.

**Supplemental Video 5:** Spline surface plots were generated to visualize spontaneous electrical signal propagation in the 8×8 MEA grid. The video shows 283 consecutive beats obtained from a siPC1 monolayer treated with etoposide for 2 h. The time scale is shown on the right side, ranging from 0 to 5 ms, using the viridis color scale.

